# Structure of the activated Roq1 resistosome directly recognizing the pathogen effector XopQ

**DOI:** 10.1101/2020.08.13.246413

**Authors:** Raoul Martin, Tiancong Qi, Haibo Zhang, Furong Liu, Miles King, Claire Toth, Eva Nogales, Brian J. Staskawicz

## Abstract

Plants and animals detect pathogen infection via intracellular nucleotide-binding leucine-rich repeat receptors (NLRs) that directly or indirectly recognize pathogen effectors and activate an immune response. How effector sensing triggers NLR activation remains poorly understood. Here we describe the 3.8 Å resolution cryo-electron microscopy structure of the activated Roq1, an NLR native to *Nicotiana benthamiana* with a Toll-like interleukin-1 receptor (TIR) domain, bound to the *Xanthomonas* effector XopQ. Roq1 directly binds to both the predicted active site and surface residues of XopQ while forming a tetrameric resistosome that brings together the TIR domains for downstream immune signaling. Our results suggest a mechanism for the direct recognition of effectors by NLRs leading to the oligomerization-dependent activation of a plant resistosome and signaling by the TIR domain.

**One Sentence Summary:** Visualization of an activated plant immune receptor that triggers the immune response upon pathogen recognition.

---

Plants have a sophisticated and finely tuned innate immune system that recognizes invading phytopathogens to protect from infection and disease. Pathogen recognition is facilitated by both membrane-anchored pattern recognition receptors and intracellular innate immune receptors (*1*). The latter include the nucleotide binding leucine-rich repeat receptors (NLRs) (*2*). While some NLR immune receptors directly bind pathogen effector proteins, others monitor effector-mediated alterations of host targets to activate effector-triggered immunity (ETI) (*3*–*8*). ETI activation is often accompanied by localized cell death referred to as the hypersensitive response (HR). Animals also employ NLR proteins as intracellular immune receptors to recognize potential pathogens and the NLR domain architecture is highly conserved, with each region playing a specific role in its mechanism of action (*9*). Plant NLRs generally consist of three domains: an N-terminal region that is either a coiled-coil (CC) domain or a Toll/interleukin 1 receptor (TIR) domain, a central nucleotide binding NB-ARC domain, and the C-terminal leucine-rich repeat (LRR) domain (*2*). NLRs are divided into TIR-NLRs (TNLs), CC-NLRs (CNLs) and CC_R_-NLRs (RNLs) based on their N-terminal domains, with experimental evidence consistently suggesting that oligomerization of the N-terminal domains is required for signal transduction and ETI activation (*3*).

While the activation mechanism of a plant CC-NLR-type resistosome has been elucidated recently (*10*, *11*), the mechanism of TNL activation remains elusive. There is still no structural evidence for TNL resistosome formation. Recently, TIR domains of both plant and animal NLRs were reported to possess an NADase activity that requires TIR domain oligomerization and leads to immune response activation (*12*, *13*). Whether the NADase activity of TIR domain is fully responsible for ETI activation, and why NAD+ cleaving only happens in the presence of TIR self-association requires further investigation.

To further our understanding of the molecular events that control the direct recognition of pathogen effectors and activation of TNL immune receptors, we solved a high-resolution cryo-EM structure of the TNL Roq1 bound to the *Xanthomonas euvesicatoria* type III effector XopQ. The Roq1-XopQ complex was directly purified from the leaves of Roq1’s native host, *Nicotiana benthamiana.* Recognition of XopQ, as well as other effectors, by Roq1 has been shown to trigger downstream ETI signal transduction, leading to a hypersensitive cell death response and resistance to pathogen invasion (*14*–*17*). Our structural data reveal that Roq1 directly binds XopQ to activate a tetrameric resistosome. We identify a series of necessary contacts for XopQ recognition by Roq1 and describe the structure of a previously uncharacterized Post-LRR domain (PL) that is essential in effector binding. We also describe the overall oligomeric state of Roq1 and the interfaces formed by the NB-ARC domain in an open conformation. Finally, we provide an explanation for the requirements of oligomerization in TIR activation, involving opening of the NADase active site in an interface-dependent manner. Together, our results provide a structural basis for direct effector recognition, oligomerization and activation of TNLs that is essential in understanding how these immune receptors detect pathogens and signal an immune response.

## Overall Structure of the Roq1 resistosome

XopQ recognition by Roq1 triggers a rapid cell death response in wild type *N. benthamiana* leaves, making it difficult to obtain sufficient protein for expression and purification (*14*). All plant TNLs require the downstream EDS1 protein to achieve cell death (*18*). In order to obtain live tissue for protein purification, we transiently co-expressed Roq1 and XopQ by Agrobacterium-mediated transformation in CRISPR-induced *eds1* mutants of *N. benthamiana* that is known to prevent Roq1-induced cell death (*15*). Cryo-EM imaging and analysis of the affinity-purified complex yielded a reconstruction at 3.8 Å overall resolution with C4 symmetry imposed (Fig. S1,S2) and showed that the Roq1 protomers assemble into a tetrameric, four-leaf clover structure with XopQ present in a 1:1 ratio (Fig. 1). Further image processing was required in order to allow building of atomic models (Fig. S3). We found that the NBD-HD1-WHD domains provide the necessary contacts for Roq1 oligomerization and bring together the four TIR domains. The LRR features the characteristic horseshoe shape and wraps around the XopQ effector protein, recognizing its surface residues. The cryo-EM map also reveals a previously uncharacterized Post-LRR (PL) domain at the C-terminal end of the LRR connected by a short nine residue linker (Fig. 1A, 2). In order to improve the density of Roq1 bound to XopQ, we applied symmetry expansion and focused refinement around the LRR-PL-XopQ region (Fig. S1,S3). The improved reconstruction showed that the XopQ effector is in its open conformation, exposing the cleft of the predicted nucleoside hydrolase active site (Fig. 2E). XopQ’s specific substrate remains unidentified, but previous studies have shown that XopQ binds ADPR, an important immune signaling molecule in plants, consistent with its immunosuppressive function (*19*).

**Fig. 1:**
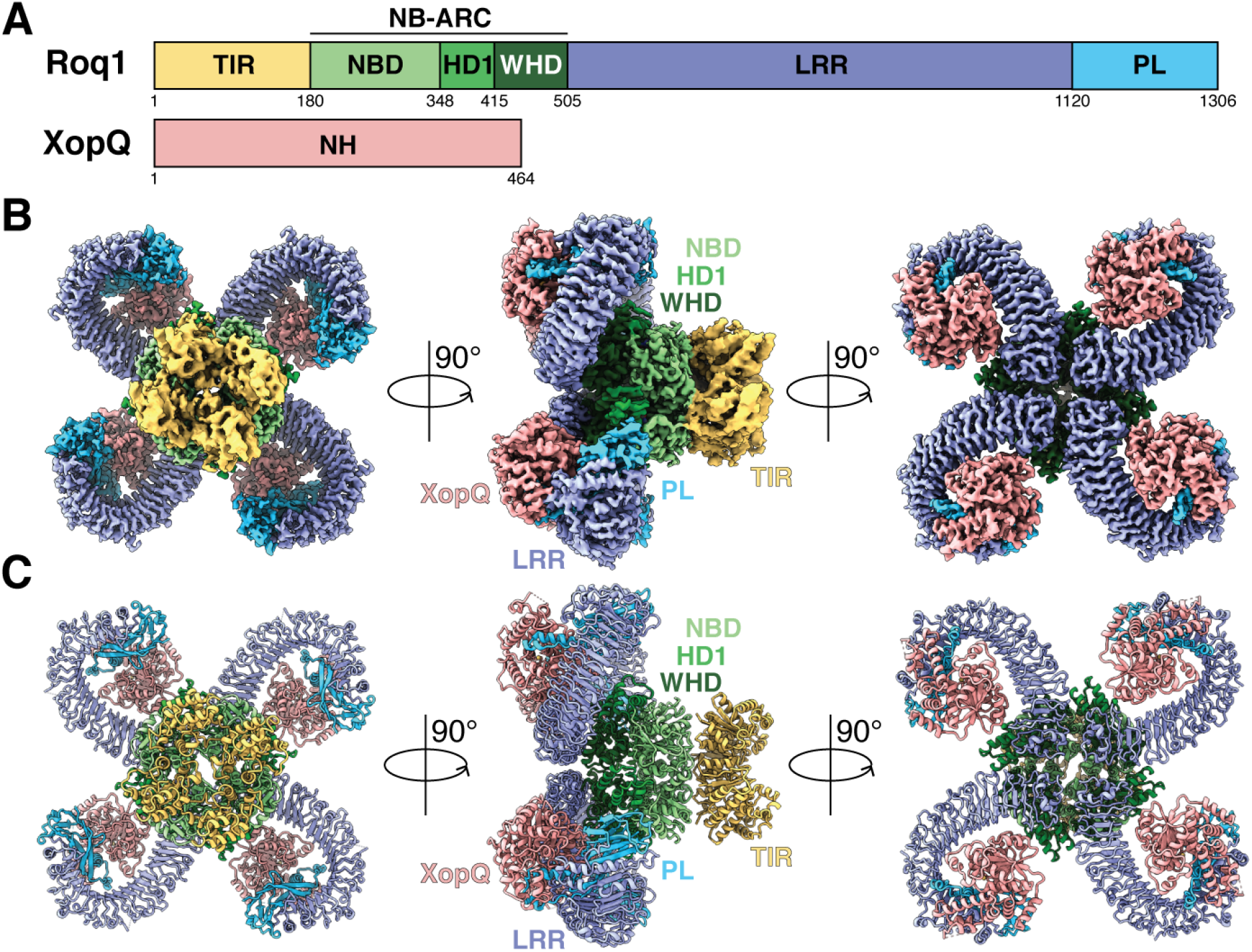
Overall structure of the Roq1-XopQ complex. (A) Schematic representations of Roq1 and XopQ with color-coded domain architecture: TIR (yellow), NB-ARC (NDB-HD1-WHD) (light green, green and dark green respectively), LRR (violet) and PL (light blue) domains, and XopQ (salmon). (B) Composite density map of the Roq1-XopQ complex from three cryo-EM reconstructions and the corresponding atomic model (C) shown in three orthogonal views. Colors according to the nomenclature in (A).

**Fig. 2:**
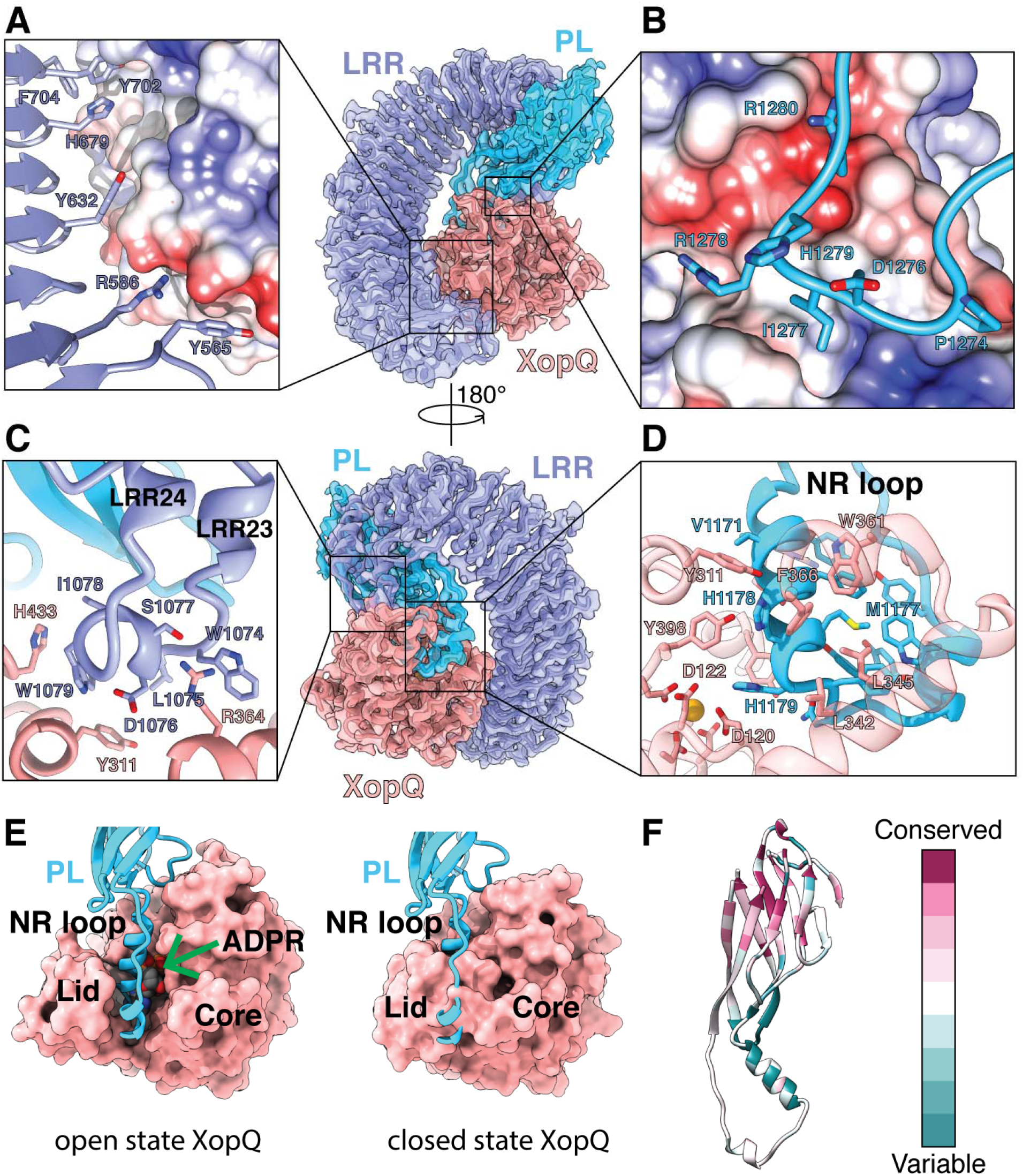
Structure of the LRR-PL region binding to XopQ. (A) Surface contacts between the N-terminal region of the LRR, shown with violet ribbon and XopQ represented by its Coulombic surface potential. (B) Surface contacts made by the loop between β-strands 7 and 8 of the PL domain shown in light blue ribbon and XopQ represented by its Coulombic surface potential. (C) The elongated LRR between repeats 23 and 24 (violet) interacting with XopQ (salmon). (D) Interactions between the NR loop (light blue) and active site residues of XopQ required for ADPR-binding (salmon). Catalytic Ca^2+^ shown in gold. (E) Left: Structure of XopQ (salmon) in the open conformation built from our cryo-EM density, with the NR loop (light blue) inserted into the active site cleft. The position of ADPR (green arrow) from the close state of XopQ (PDB: 4P5F) is modeled to show its overlapping position with the NR loop. Right: The ADPR-bound, closed state of XopQ (PDB: 4P5F). The NR loop is modeled to demonstrate the clashes that would occur upon XopQ closure. (F) Residue conservation of the PL domain calculated in Consurf (*52*). Some residues had unreliable conservation scores due to insufficient data and are represented in white, corresponding to a neutral score.

## Recognition of XopQ by Roq1

The 24 leucine-rich repeats of Roq1 form a 150 Å long scaffold that bends around XopQ, displaying key contact residues along its surface (Fig. 2). We find the LRR of Roq1 interacts with the effector in two different ways: (i) In the region where XopQ is in close contact with the LRR scaffold, several side chains exposed on the surface of the LRR directly interact with the substrate (Figure 2,A). A similar mechanism is utilized by the LRRs of the CC-NLR ZAR1 to recognize RKS1 and TLR3 to recognize dsRNA (*11*, *20*). The majority of these residues have large aromatic side chains that recognize hydrophobic patches and grooves on the surface of XopQ. (ii) In regions where the LRR scaffold is too far away to interact with the effector directly, we find an elongated linker between two leucine-rich repeats (LRRs 23 and 24) that reaches over to bind XopQ (Fig. 2C). A small amphipathic α-helix is formed at the site of contact, with hydrophobic side-chains recognizing conserved residues at the outer edge of XopQ’s active site cleft (Y311^XopQ^, H433^XopQ^) (Fig. 2C). The extended linker then loops back towards the scaffold and forms the next repeat in the LRR. Mutating the residues that form the hydrophobic face of the α-helix (L1075^Roq1^, W1079^Roq1^, I1078^Roq1^) to alanines resulted in loss of the HR phenotype, suggesting these interactions are critical for XopQ recognition (Fig. S4).

A 9-residue linker (aa. 1120-1129) connects the C-terminal end of the LRR to the Post-LRR (PL) domain, which also interacts with XopQ (Fig 2B,D). This domain folds into a β-sandwich, with 9 antiparallel β-strands arranged in two β-sheets (Fig. S5). The last LRR forms hydrogen bonds with one of the β-strands, thereby rigidifying the conformation between the two domains. Because PLs serving in pathogen detection have been found in other TNLs, but remain poorly characterized, we sought to further investigate possible structural homology of the Roq1-PL domain with published structures (*21*). Analysis using the CATH database (*22*) revealed proteins with immunoglobulin-like folds or jelly-roll folds as the closest structural homologues. Strikingly, the mode of recognition utilized by the PL domain to detect the foreign protein is reminiscent of the way immunoglobulins bind to their antigen (Fig. S6). Loops emerging from the β-sandwich structure target sites in XopQ to form substrate-specific contacts with the pathogen protein in a manner that resembles the complementarity-determining regions found in antibodies. The loop between β-strands 7 and 8 of the PL domain simultaneously recognizes a hydrophobic pocket via the insertion of an isoleucine (I1277^Roq1^), and an area of negative potential targeted by R1280^Roq1^ (Fig. 2B). Disrupting these interactions with a Roq1 double mutant (R1280D and I1277A) prevented HR in tobacco leaves, suggesting that these interactions are essential for XopQ detection by Roq1 (Fig. S4).

The greatest number of contacts between the PL domain and XopQ are made by a 33 residue loop (aa 1163-1196) that dives into the active site cleft of the effector and positions sidechains in close contact with conserved sites required for ADPR-binding (Fig. 2D,E) (*19*). We refer to this loop as the NR loop for its ability to bind residues in XopQ responsible for nucleoside recognition (NR). Two α-helical segments of the loop bring together large hydrophobic sidechains that interface with the interior lid region of XopQ (Fig. 2D,E). The conserved XopQ residues targeted in this region (W361^XopQ^, F366^XopQ^, L345^XopQ^) serve to recognize the base moiety of ADPR (*19*). Active site residues that would otherwise stabilize the α-phosphate of the ligand (Y311^XopQ^ and Y398^XopQ^) are recognized by Roq1 V1171^Roq1^, H1178 ^Roq1^ and H1179^Roq1^ (*19*). Additionally, H1179^Roq1^ interacts with D120^XopQ^ which is involved in the recognition of one of the sugar moieties in ADPR (*19*). In summary, the conserved residues in XopQ involved in recognizing the base, α-phosphate and ribose moieties in ADPR are targeted by the Roq1’s NR loop. Mutating the NR loop to a short flexible linker (-SGGGSGGS-) resulted in loss of HR, suggesting the Roq1 mutant could no longer recognize XopQ (Fig. S4). Comparison of our structure of XopQ, which is in an open state, with the closed, ADPR-bound state (PDB: 4P5F), shows that the NR loop overlaps with ADPR and thus would prevent the ligand from entering the active site cleft or interacting with XopQ (Fig. 2E). The presence of the NR loop may also block XopQ from transitioning to the closed state, as the NR loop would clash with the lid region capping the active site. These observations lead us to hypothesize that Roq1 not only recognizes the pathogen but may also inhibits the virulence function of XopQ (*23*).

The newly discovered PL domain of Roq1 has a conserved β-sandwich core that may be found in the NLRs of other members of the nightshade family (Figure 2,F). We ran a BLAST search using the sequence of the PL domain and found multiple hits corresponding to resistance genes in other species of tobacco, as well as in various species of potatoes, peppers and morning-glories. The more conserved residues are within the strands of the β-sandwich, whereas the loop residues pointing towards XopQ are more variable (Fig. 2F). The NR loop is only found in three other tobacco species, with minor sequence differences (V1171→I, Y1195→F). This pattern of conservation suggests that the variable loops emerging from the PL domain core of related NLRs could serve in recognizing different pathogen effectors via a similar mechanism to Roq1. Such a strategy would be akin to that of sequence variations in the complementarity-determining regions of antibodies that enable them to recognize a diversity of epitopes.

Previous studies suggest that it was difficult to identify mutations in XopQ that could evade Roq1 recognition (*24*). This is consistent with our results that demonstrate that the LRR and PL domains of Roq1 make multiple contacts with XopQ, suggesting that this gene would be durable in the field and difficult for the pathogen to evade. In the future, these contacts could be modified to build synthetic receptors targeting various pathogen effectors resulting in novel recognitional specificities.

## Oligomerization of Roq1 mediated by the NB-ARC domain

NLRs are generally thought to exist in an inhibited state mediated by either intra- or intermolecular contacts that prevent oligomerization between protomers and activation of the immune response (*9*, *25*, *26*). Structural studies of inactive NLRs suggest that these inhibitory contacts hold the nucleotide-binding region (NBD, HD1, WHD) in an closed state (*10*, *27*–*30*). Upon activation, these interactions must be disrupted in order to transition to the oligomerization-prone state, were the WHD is moved away from the nucleotide-binding site, thereby displacing the ADP-specific Met–His–Asp motif on the WHD and allowing ATP or dATP-binding (NLR have been shown to bind either ATP or dATP in their active state, in this study we chose to stick with ATP for simplicity) (*11*, *31*–*35*). We expect Roq1 to be similarly regulated by autoinhibitory contacts with the LRR, based on evidence demonstrating that a truncated version of Roq1 missing the LRR-PL region spontaneously triggers an immune response in the absence of effectors (*15*). We sought to determine if the PL domain could play a role in auto-inhibition. Removing the PL domain of Roq1 (ΔPL) resulted in loss of HR *in planta*, suggesting the LRR, not the PL domain, is involved in making the intramolecular contacts that obstruct a conformational switch to the active state (Fig. S4).

Four Roq1 protomers oligomerize via the NB-ARC domains upon substrate recognition. Our density map reveals the molecular contacts between the three subdomains of the NB-ARC (NBD, HD1 and WHD) in the context of the resistosome, as well as the presence of ATP at the nucleotide binding pocket, consistent with an activated state of Roq1 (Fig. 3). The ATP molecule is stabilized at the interface between the NBD and HD1, with the NBD recognizing the β-phosphate via the canonical P-loop (K224 and T225), two aspartates (D300 and D301) of the Walker B motifs in close proximity to the ATP γ-phosphate, and additional residues recognizing the sugar and base moieties (Fig. 3A) (*26*). In agreement with published NLR structures in the active state (*11*, *32*–*35*), the WHD of activated Roq1 is rotated away from the nucleotide-binding site, thereby exposing the oligomerization interface (Fig. 3B,C). This arrangement allows the NBD-HD1 surface of a protomer to intercalate with the NBD-WHD surface of its neighbor. The major interactions involve the HD1-WHD interface and an NBD-NBD interface (Fig. 3C). HD1 binds to the neighboring WHD using a mixture of polar and hydrophobic contacts. Residues surrounding the fourth α-helix of HD1 (aa 401-413) play an important role in forming the Roq1 tetramer (Fig. 3C, left). Single point mutations changing the character of these residues (E399R, V403D and R410A) resulted in loss of HR, suggesting Roq1 oligomerization was disrupted (Fig. S4). Similar results were observed when mutating the charged residues that bring together NBD domains (R229D) (Fig. S4). In other structures of multimeric NLRs (*11*, *32*–*35*), the contacts between the NBDs also occur via their N-terminal linker. The equivalent linker in Roq1 is poorly resolved in our cryo-EM map compared with the surrounding NBD, for which we observe well-defined sidechain densities, indicating that the linker region in Roq1 is flexible in the tetrameric state. Interestingly, the same linker in other NLRs provides contacts that are in part responsible for properly positioning the NBDs relative to each other. In fact, in NLRs that require many protomers for assembly, the linker forms an α-helix, whereas in smaller complexes, such as the pentameric ZAR1, the N-terminal linker forms a slim structured loop without any secondary structure, allowing for the NBD to pack more tightly (Fig. S7). Similarly, the poorly defined structure in the Roq1 linker could explain the tight packing between NBDs that results in tetramerization instead of higher oligomeric states.

**Fig. 3:**
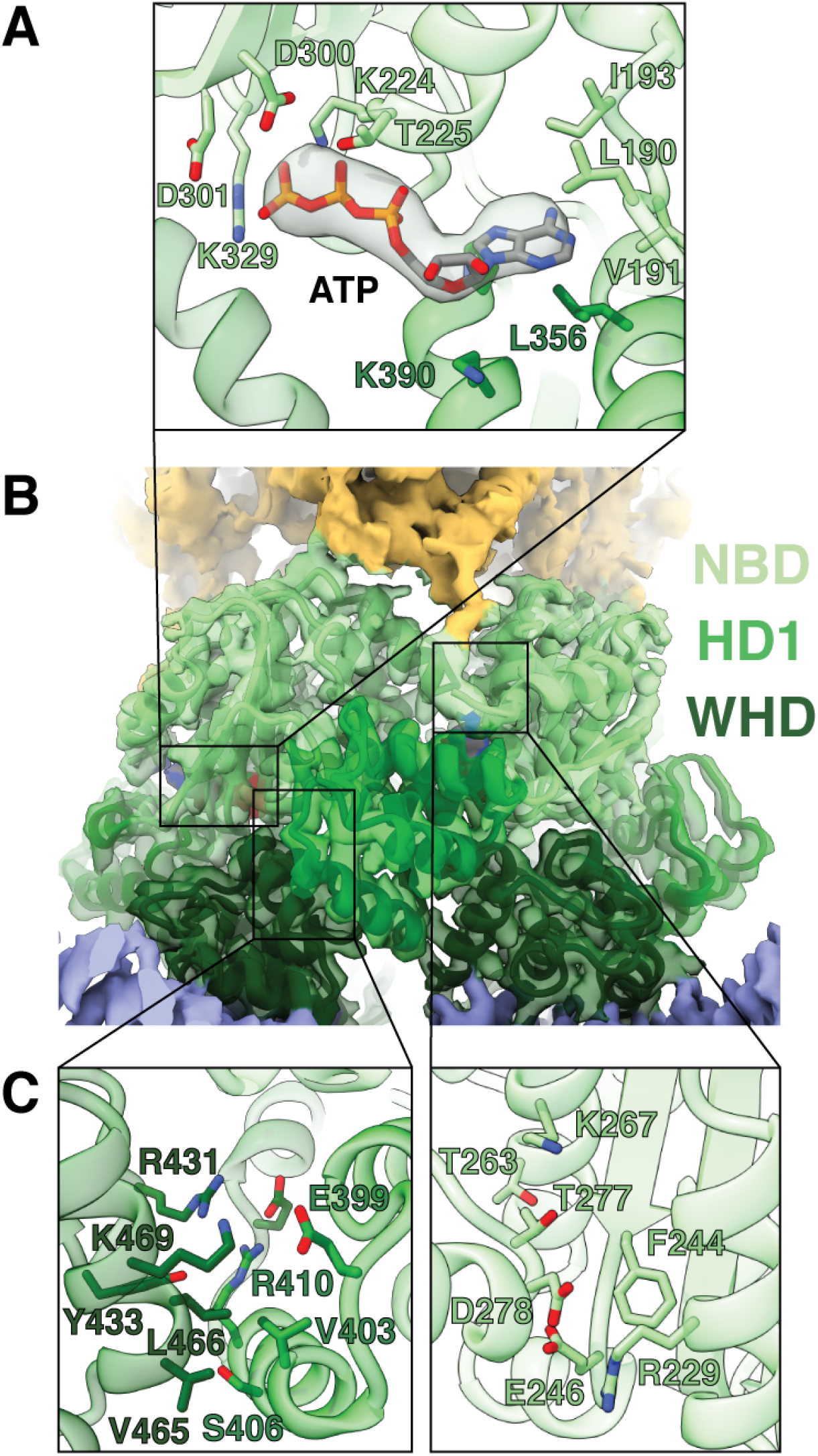
Oligomerization interfaces between NB-ARC domains. (A) ATP modeled in the cryo-EM density (gray) at oligomerization interfaces, showing the sidechains of residues involved in ATP binding. (B) Interface between two NBARC domains with the NBD (light green) and WHD (dark green) of one protomer intercalating with the NBD (light green) and the HD1 (green) of the neighboring one (C) Left: Residues involved in contacts between the WHD (dark green) and HD1 (green). Right: Residues involved in contacts between neighboring NBDs.

Mechanisms have been proposed for the oligomerization of NLRs with differing reliance on nucleotide-binding (*25*). In the case of ZAR1, indirect substrate recognition mediated by the guard protein, RKS1, causes a conformational change in the NBD and triggers ADP-release, but the individual ZAR1 protomers are still unable to oligomerize independently of ATP (*10*, *11*). In contrast, the direct recognition of flagellin by the NLR NAIP5 induces a large conformational transition to the active state (*34*, *36*) and has been shown to activate even when the ATP-binding P-loop motif was mutated (*37*). The structure of the Roq1 NB-ARC domain closely resembles that of ZAR1 (Fig. S8) and shares a 22.2% sequence similarity. Previous studies have also shown that mutation in the P-loop of Roq1 prevented oligomerization (*15*), suggesting ATP-binding is required for assembly. Based on these observations, we expect Roq1 to follow a similar oligomerization mechanism to ZAR1 in which substrate recognition by the LRR-PL of Roq1 induces a conformational change in the NBD that releases ADP. ATP-binding would then be required to transition to the oligomerization-prone state.

## TIR domain oligomerization and activation

Tetramerization of the NB-ARC domains brings the TIR domains into close proximity (Fig. 4). The individual TIR domains interact with each other upon resistosome assembly, allowing them to become active NADases and trigger HR (*38*). The mechanisms for how this association renders TIRs catalytically active remains poorly understood. Many structural studies on TNL have relied on truncated TIR-containing proteins that are missing the subunits driving oligomerization (*13*, *39*–*43*). Here, we describe a mechanism for TIR association and activation in context of the fully assembled Roq1 TNL.

**Fig 4:**
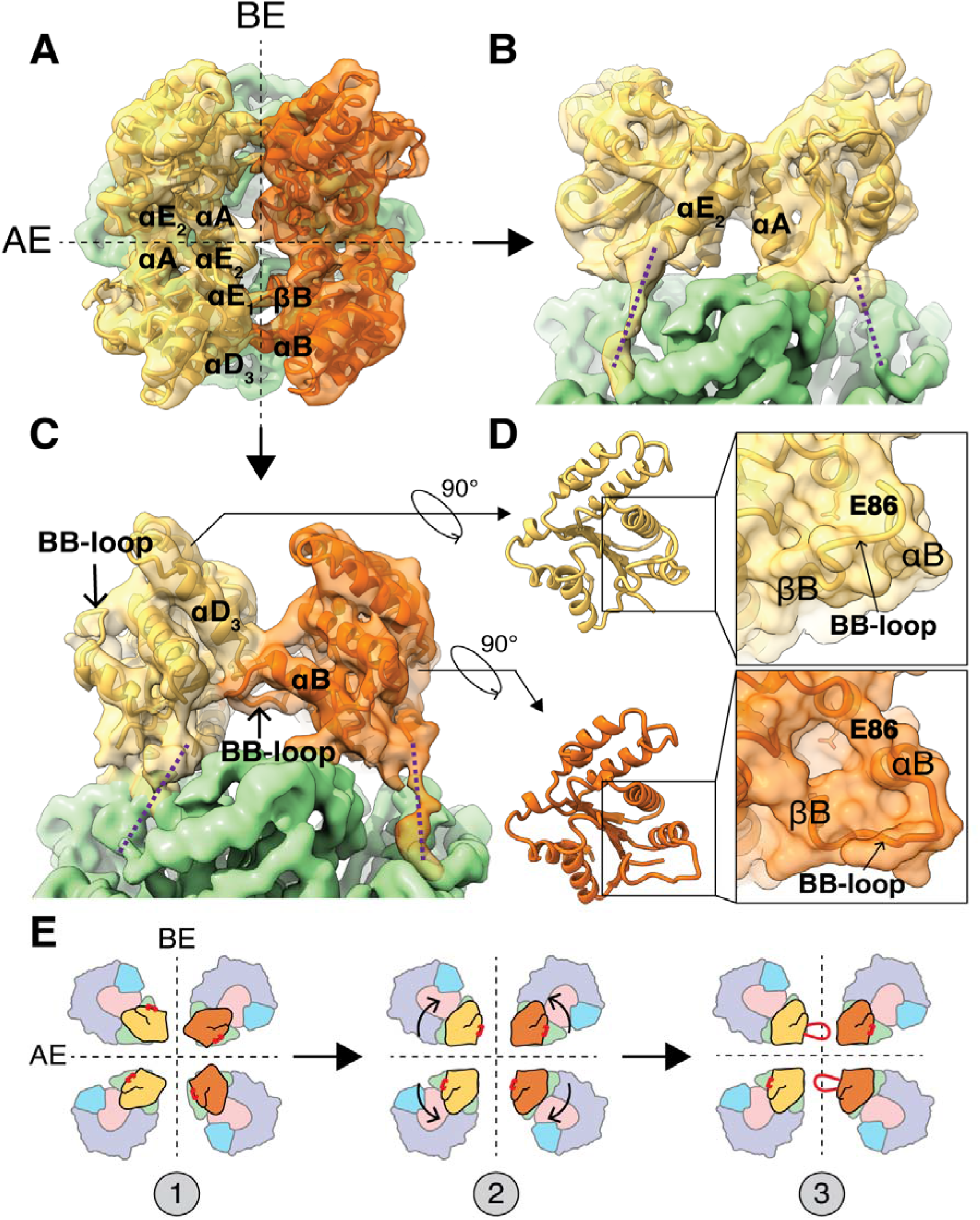
TIR domain interfaces and conformational rearrangement of the BB-loop. (A) Top view of Roq1 displaying the four TIR domains organized as a head-to-tail dimer of head-to-head dimers (each symmetric dimer shown in distinct yellow and orange). The two distinct interfaces are marked with black dotted lines: the AE interface is formed between TIR domains shown in the same color (either yellow or orange). (B) Orthogonal view from panel A of the AE interface. The proposed paths of the protein chain linking the TIR domain (yellow) to the NBD (light green) are drawn with purple dotted lines. (C) Orthogonal view from panel A of the BE interface marking the BB-loop positioned under the αD_3_ to αE_1_ helices. The proposed paths of the protein chain linking the TIR domain (yellow and orange) to the NBD (light green) are drawn with purple dotted lines. (D) Top: NADase active site of a TIR domain for which the BB-loop is not interfacing with the DE surface. Bottom: Conformational rearrangement in the BB-loop bound to the DE-surface. The side chain of the catalytic glutamate (E86) is shown in stick representation. (E) Hypothetical mechanism of TIR oligomerization with the position of the BB-loop in red. (1) Individual TIR domain from four Roq1 protomers are brought in close proximity. (2) TIR domains recognize each other at the AE and BE interface. (3) Assembly causes the conformational rearrangement in the BB-loop that opens the NADase active site.

Our initial four-fold symmetric reconstruction of Roq1-XopQ could not clearly resolve the density corresponding to the TIR domains. Further analysis (see Methods) revealed that the four TIR domains do not assemble in a four-fold symmetric fashion but form an only two-fold symmetric dimer of dimers. The change in symmetry at the TIR domains highlights the importance for flexibility in the linker that connects them to the NBDs, as discussed previously. After adjusting the symmetry for this region and focused refinement, the TIR domains reached an overall resolution of 4.6 Å, allowing us to visualize secondary structure elements and trace the polypeptide backbone (Fig. S1,3). The TIR domains are arranged forming two types of interfaces. First, TIR domains engage in a head-to-head, symmetric interaction involving alpha helices the αA and αE_2_ of each protomer (nomenclature of TIR structural motives as in (*44*)) (Fig, 4A,B, shown as interaction between same color protomers). This interface, previously termed AE-interface, is also found in many crystal structures of isolated plant TIR domains, including RPP1, RPV1 and SNC1 (Fig. S9) (*13*, *39*–*41*, *43*). Consistent with published studies on these plant TNLs, mutating residues in the αA helix of Roq1 (H30A) disrupts HR and highlights the functional importance of these contacts (Fig. S4).

In our structure of Roq1-XopQ, TIR domains engaged in an AE-AE interaction then further dimerize head-to-tail forming what is described as the BE interface (*45*) (Figure 4A,C shown as interactions between different color protomers). In the BE interaction the so-called BB-loop (residues between βB and αB) of one TIR domain plugs beneath the loop between αD_3_ and αE_1_ of the adjacent one (Fig. 4C). Previous mutational analyses already demonstrated a functionally important role for the BB-loop in TIR domains (*13*, *46*, *47*). We further mutated residues in the αD_3_-αE_1_ loop (I151A and G153A) that are in close contact with the BB-loop and found that they independently resulted in loss of the HR phenotype (Fig. S4).

Association between plant TIR domains at the DE surface (formed in part by the αD_3_-αE_1_ loop) has previously been observed in crystallographic studies but their conformations are different from the ones defined by our cryo-EM analysis. For example, the TIR domains of RPP1, SNC1 and L6 face each other head-to-head at their DE surfaces, with different rotational angles relative to each other, instead of interacting in a head-to-tail fashion (*39*); potentially because the TIR domains were visualized in isolation and the NB-ARC domain responsible for driving oligomerization was truncated. These studies highlighted the importance of the DE surface in plant TIR domain oligomerization, but the proper interactions remained unclear due to the variability in conformation between structures. Interestingly, there is a similar BE-interaction in more distant phyla. The crystal structures of TIR domains from the human SARM1 (*13*) and MAL (*48*) proteins, as well as TRR-2 (unpublished, PDB: 4W8G and 4W8H) from *Hydra magnipapillata* share a strikingly similar BB-loop conformation to that found in the activated Roq1 tetramer, in which it fits under the αD_3_-αE_1_ loop (Fig. S10). This structural relationship suggests shared mechanistic features for TIR domain assembly and activation between animals and plants. In fact, the human SARM1 TIR domains simultaneously form AE and BE interfaces in the crystal lattice (Fig. S9,S10) (*13*).

Our structure now demonstrates that the BB-loop takes on two different conformations within the activated Roq1 tetramer and undergoes a conformational switch as TIR domains interact at a BE interface. In the protomer in which the BB-loop is unbound (the one that is contributing to the BE contacts through its DE surface), it is seen in an upward position along the rim of the NADase active site (Fig. 4C). This conformation is the same as found in the TIR crystal structures lacking BE contacts (*39*, *40*, *42*, *43*). In the other protomer, the BB-loop interfacing with the adjacent DE surface has been repositioned via a downward motion of about 12 Å (Fig. 4D, bottom). A highly conserved glycine residue (G52^Roq1^) in the BB-loop likely provides the flexibility required to undergo this conformational switch (Fig. S11). Mutating the equivalent glycine (Fig. S11) to a proline in SARM1 (G601P) was shown to hinder flexibility and prevent a transition to the engaged state, resulting in a defective BE-interface with the loop stuck in the upward position and a severe decrease in NADase activity (*13*).

Interestingly, repositioning of the BB-loop induced by the BE-interface opens the NADase active site (Fig 4D). Large, positively charged sidechains (K50^Roq1^, R51^Roq1^ and K53^Roq1^) that would otherwise crowd the entrance of the active site are moved down with the BB-loop. The structure of a SARM1 mutant (G601P), in which the BB-loop is trapped in the unengaged state, reveals a lysine inserted inside the active site cleft; this indicates these sidechains may act to prevent substrate binding (*13*). Furthermore, NADase activity increased when equivalent BB-loop arginines were mutated to alanines in the plant RUN1 TIR domain (*13*). Together, these studies suggest that these large positively charged sidechains serve to inhibit the NADase function and must be displaced for TIR activation.

Freeing the active site exposes conserved residues that have been proposed to recognize NAD+ based on biochemical and structural studies using chemical analogues (*12*, *13*). The nicotinamide moiety of the substrate is supposed to fit in the active site cleft of the TIR domain, bringing the covalently linked ribose in close proximity to the catalytic glutamate. Mutating the catalytic glutamate in Roq1 (E86^Roq1^) to an alanine abrogates HR (Fig. S4), suggesting loss of NADase activity. Mechanistic details of NAD+ cleavage and product formation remain unresolved and have been found to vary among TIR domains (*38*). The steps in this enzymatic reaction involve breaking the glycosidic bond that connects nicotinamide to ADPR and, in some cases, a structural rearrangement in ADPR that leads to the formation of cyclic-ADPR or variant-cyclic-ADPR (*12*, *49*). These products have been shown to modulate Ca^2+^ level in plant cells, which is a widely used chemical signal for responding to various biotic and abiotic stresses (*50*, *51*).

In the case of the fully assembled Roq1 TNL, it is clear that the AE and BE interfaces are essential in TIR signaling. Both interfaces align the TIR domain in a conformation inducive to NADase active site opening (Fig. 4E). Whether this mechanism of TIR association can be applied to other TNL remains to be determined. There likely exist alternative ways for TIR domain to assemble, based on the number of protomers required to build the active complex, hetero-complex formation with other NLRs, and interface requirement for activation.

## Summary

Our structure of the Roq1-XopQ complex, together with previous biochemical studies, lead us to propose a mechanism for TNL immune signaling: (i) both the LRR and PL domains recognize the pathogen effector, at which point the NR loop inserts itself into the active site cleft of XopQ and targets conserved residues required for nucleoside-binding; (ii) the NB-ARC domain is released by the LRR and undergoes a conformational switch to the ATP-bound oligomerization state; (iii) Roq1 protomers associate into a four-leaf clover structure and the TIR domains are brought in close contact; (iv) the TIR domains bind to each other forming distinct AE and BE interfaces and causing a conformational rearrangement in the BB-loop of two of the subunits; (v) the NADase active site is exposed, allowing for the cleavage of NAD+. Future work will also be required to identify the exact molecular species produced by the TIR domains and how they are utilized by the immune system of the host to trigger a response to pathogen invasion.

## Supporting information

Movie S1: Cryo-EM density map and atomic model of the Roq1-XopQ complex with colors corresponding to the different protein domains (as in Fig. 1).

## Acknowledgments

We thank Patricia Grob and Daniel Toso for electron microscopy support, and Abhiram Chintangal and Paul Tobias for computational support. Data was collected at the BACEM facility with the Berkeley QB3. We thank Nicole Haloupek and Patricia Grob for advice with making cryo-EM grids of NLR samples. We thank Basil Greber for advice with data processing and with model building and refinement. We thank Ksenia Krasileva, Daniil Prigozhin, and Kirsten Verster for useful discussion in regards to PL domain conservation and LRR-effector interactions. We thank Bostjan Kobe for useful discussion on TIR domains. Molecular graphics and analyses performed with UCSF Chimera and UCSF ChimeraX.

## Author contributions

Work was supervised by E.N. and B.J.S. T.Q. constructed Roq1 and XopQ vectors, expressed the Roq1-XopQ protein complex in *N. benthamiana*. R.M. and T.Q. established purification protocols and purified the Roq1-XopQ complex. R.M. prepared cryo-EM grids, collected and processed cryo-EM data, and carried out model building, refinement and interpretation. C.T assisted in purification and grid making. T.Q., H.Z., R.M. and F.L. constructed Roq1 mutants. T.Q. and H.Z. detected protein expressions and HR phenotypes. M.K. assisted in Roq1 and XopQ protein expression in *N. benthamiana*. R.M. wrote the initial draft of the manuscript. All authors contributed to the final version of the paper.

## Data and materials availability

CryoEM density maps and fitted models have been deposited in the Electron Microscopy Data Bank (EMDB) and the Protein Data Bank (PDB). The maps for initial reconstruction of the Roq1-XopQ complex, focused refinement around the LRR-PL-XopQ region and TIR domains have been deposited with the following EMDB ascension codes 22381, 22380 and 22383. The refined coordinate models have been deposited with the following PDB ascension codes 7JLV, 7JLU and 7JLX respectively.

## Supplementary Materials

### Materials and Methods

#### Plant Materials and Growth Conditions

The N. benthamiana eds1-1 mutant was described as previously (14). The binary vector containing Roq1 guide sequence (GATGATAAGGAGTTAAAGAG) and Cas9 was described previously (15) was transformed into agrobacterium and used for generating N. tabacum roq1-1 stable mutant plants by CRISPR-Cas9 gene editing system. N. benthamiana and N. tabacum plants were grown in a growth chamber under a 8-hr-light/16-hr-dark photoperiod at 23-25□.

#### Expression and Purification of the Roq1-XopQ Complex

Roq1 and XopQ were fused with C-terminal 3xFlag tag and N-terminal StrepII tag, respectively, and transformed into Agrobacterium GV3101. The agrobacterium GV3101 strains containing Roq1-3xFlag and StrepII-XopQ were co-inoculated into N. benthamiana eds1 mutant leaves. At 30 hours after infiltration, 200g of leaves were harvested and ground using a mortar and pestle and resuspended in 400 mL of Lysis Buffer (50 mM HEPES pH 7.5, 1 mM EDTA. 5 mM MgCl2, 150 mM NaCl, 10 mM KCl, 0.4% NP40, 5% glycerol, 10 mM DTT). Leaves were further lysed by sonication for 2 min at 20 kHz. The cell lysate was initially centrifuged at 18,000xg for 45 min to pellet large debris and the harvested supernatant was further centrifuged at 40,000xg for 45 min to remove any smaller residuals. The clarified extract was then incubated with an ANTI-FLAG M2 affinity gel (Sigma-Aldrich) for 3 hours at 4 ◻C. The gel was washed with Wash Buffer (20 mM HEPES pH 7.5, 1mM EDTA, 5 mM MgCl2, 150 mM NaCl, 10 mM KCl, 0.2% NP-40, 10% glycerol) and the sample was eluted twice by incubating in Wash Buffer supplemented with 300 μg/mL of 3xFlag peptide for 30 min. Eluates were pooled and incubated with Strep-Tactin Superflow Plus (Qiagen) resin for 1 hour. The resin was then washed with Wash Buffer and the totality of the sample was eluted in 5 sequential steps by adding Wash Buffer supplemented with 10 mM Biotin. The protein complex was flash frozen in liquid N2 and stored at −80°C. Individual steps can be visualized by SDS-PAGE in Fig. S12.

#### Sample Preparation for Cryo-EM

QUANTIFOIL R2/2 holey carbon grids were coated with a thin film of continuous carbon and plasma cleaned (Tergeo-EM, PIE Scientific LLC) before addition of sample. Because of the low concentration of the Roq1-XopQ complex in our sample, we attempted to float the carbon-coated grid on the sample drop to enable prolonged incubation to allow attachment of more molecules to the carbon. However, due to the high detergent concentration of the sample buffer, loss of surface tension caused the grids to sink to the bottom of the drop. Therefore, we used a Teflon well to hold 20 μL of sample (supplemented with 1 mM ATP) and deposited the grid in the well carbon side up, upon which the grid fell to the bottom of the well, allowing sample adsorption to the carbon-coated side. The grid was incubated with the sample for 90 min at 4° C. It was then removed from the drop and washed in a 50 μL drop of cryo-EM-friendly buffer (10 mM HEPES pH 7.5, 1 mM EDTA, 5 mM MgCl_2_, 150 mM NaCl, 10 mM KCl, 3% trehalose). We gently blotted the grid using filter paper and added 4 μL of cryo-EM-friendly Buffer before mounting the grid onto a Thermo Fisher Scientific Vitrobot Mk. IV. The grid was immediately blotted for 10 sec (blot force 10) and plunge-frozen in liquid ethane.

#### Data Collection

The grid was loaded onto a Titan Krios cryo-electron microscope (Thermo Fisher Scientific) operating at 300 kV and equipped with a K3 direct electron detector camera (Gatan) mounted behind a BioQuantum energy filter (Gatan). Electron micrographs were acquired as dose-fractionated movies (11,134 movies in total) in super-resolution counting mode with the microscope set to 53,271x magnification (corresponding to a pixel size of 0.9386 Å) and a total electron exposure of 50 e^−^/Å^2^. Defocus values ranged from −0.9 to −2.5 μm. Automated data collection was controlled by SerialEM. For high-throughput data collection, we used image shift with active beam tilt correction enabled to collect 20 movies at each stage position. All other parameters can be found in Table S1.

Expression and Purification of the Roq1-XopQ Complex

#### Data Collection

The grid was loaded onto a Titan Krios cryo-electron microscope (Thermo Fisher Scientific) operating at 300 kV and equipped with a K3 direct electron detector camera (Gatan) mounted behind a BioQuantum energy filter (Gatan). Electron micrographs were acquired as dose-fractionated movies (11,134 movies in total) in super-resolution counting mode with the microscope set to 53,271x magnification (corresponding to a pixel size of 0.9386 Å) and a total electron exposure of 50 e^−^/Å^2^. Defocus values ranged from −0.9 to −2.5 μm. Automated data collection was controlled by SerialEM. For high-throughput data collection, we used image shift with active beam tilt correction enabled to collect 20 movies at each stage position. All other parameters can be found in Table S1.

#### Data Processing

All processing steps were performed using RELION 3.1 (*53*) unless otherwise indicated. Movies were imported into RELION and classified into 9 optics groups according to the respective beam shift used during acquisition. Alignment of the movie frames was performed using MotionCor2 (*54*) and GCTF (*55*) was used for fitting of the contrast transfer function and defocus estimation. To ensure that we captured particles in all poses present on the grid, we used the unbiased Laplacian-of-Gaussian autopicker (*56*) in RELION for particle picking. Instead of 2D classification, an initial 3D classification (with C4 symmetry applied) was performed in order to prevent loss of rare views that might be classified into classes containing broken particles or false-positive particle picks in 2D classification. An initial reconstruction of the Roq1-XopQ complex generated in cryoSPARC (*57*) from a grid quality screening session was used as the reference model. The particles from the best classes in this initial 3D classification were subjected to successive rounds of alignment-free 3D classification, and alignment-free 2D classification for each 3D class, followed by removal of bad particles and 3D refinement. This enabled us to recover the side views of the Roq1-XopQ complex, which we failed to do using alternative processing approaches. A final round of 3D-refinement and alignment-free 3D classification (tau = 16) yielded one high-quality class containing 15,263 particles, with a broad distribution of projection directions. After CTF-refinement and Bayesian polishing (*56*) of these particles, 3D-refinement resulted in a reconstruction of the Roq1-XopQ complex at 3.8 Å resolution overall (FSC = 0.143). This initial map was of sufficient quality for atomic model building of the NB-ARC region (NBD-HD1-WHD) as well as the most N-terminal portion of the LRR. Further processing was needed to improve the LRR-PL-XopQ and TIR domain regions.

To resolve the interaction between Roq1 and XopQ, we applied symmetry expansion to our particles and performed a focused refinement using a mask around one LRR-PL-XopQ module. The angular search space was restricted to preserve the particle orientations after symmetry expansion. The resulting reconstruction converged to 3.8 Å resolution, displaying clear separation between β-strand of the LRR and PL domain, as well as densities for the sidechains that interact with XopQ.

To improve the TIR domain map, we applied signal subtraction to our particles using a 3D mask around the TIR domains and selected particles exhibiting good density in this region by alignment-free 3D classification. This improved the overall signal for the TIR domain but the features of the density were too poor to confidently fit a model into this four-fold symmetric map. We reasoned that the poor map quality might originate from a symmetry mismatch between the TIR domains and the remainder of the complex, with the TIR domains possibly assuming lower symmetry. Therefore, we applied symmetry expansion to the particles subset and classified the data using alignment-free classification after signal subtraction to remove everything except the TIR domains from the particle images. The particles split equally into two identical classes rotated 90° relative to each other, revealing four TIR domains forming a dimer of dimers with C2 symmetry. Focused refinement of the TIR and NB-ARC regions (the signal from TIR domains alone was too small for proper particle alignment) improved the overall resolution to 4.6 Å, but the NB-ARC region is better resolved than the TIR domains. Based on local resolution estimation, the resolution of the TIR domains is around 7.5 Å, with the highest resolution features observed at the interface between the TIR domains.

#### Model Building and Refinement

Each of the three cryo-EM maps (NB-ARC region, LRR-PL-XopQ module, and TIR domains) were used separately to build atomic models of the different parts of the Roq1-XopQ complex. Initially, a model generated using SWISS-MODEL (*58*) based on the structure of ZAR1 NBD (PDB: 6J5T) was docked into our cryo-EM map using UCSF Chimera. This model served as a starting point to build the structure of the NB-ARC domain manually in COOT (*59*). Our initial map of the Roq1-XopQ complex was used to build residues 189-625 of Roq1 as well as for fitting the ATP ligand. The map resulting from the focused classification around the LRR-PL-XopQ region was then used to build the following C-terminal residues of Roq1. We used the secondary-structure prediction algorithm in Phyre2 (*60*) to guide us in building our model, specifically matching densities of β-strands and α-helices to sequences of residues with a corresponding predicted secondary structure. In a few cases, poorly resolved linker regions between β-strands were left unmodeled. We used the densities of large side chains and nearby secondary structure elements to ensure correct register assignment of the residues that followed. The structure of the open conformation of XopQ from *Xanthomonas oryzae pv. oryzae* (PDB: 4KL0) was used to fit XopQ into our density. The minor sequence differences with XopQ from *Xanthomonas euvesicatoria* were fixed in COOT (*59*).

A model for the Roq1 TIR domain was generated in SWISS-MODEL (*58*) using the structure of the plant RPV1 TIR domain (PDB: 5KU7), as it shares the most sequence similarity among published structures. Four individual TIR domain monomers were docked in the TIR domain map using Chimera (*61*) and modified in COOT (*59*) to properly fit the density.

The atomic models for the TIR domain, the NB-ARC domain and LRR-PL-XopQ region were each refined in their respective maps with successive rounds of real-space refinement in PHENIX (*62*). Ramachandran, rotamer, C_b_, and secondary structure restraints were applied throughout to maintain good model geometry at the resolution of our cryo-EM maps. Structural issues were corrected manually in COOT (*59*) between rounds of refinement. The model was validated using MOLPROBITY (*63*) within PHENIX (*62*), and the model vs. map FSC was calculated using the MTRIAGE (*64*) validation tool in PHENIX (*62*).

#### HR Phenotype and Protein Expression

The various Roq1 mutants were made and constructed into PE1776 vectors fusing with a C-terminal 3Flag tag. For HR phenotype observation, Roq1 mutants and StrepII-XopQ were transiently co-expressed in *Nicotiana tabacum roq1-1* mutant leaves via agrobacterium mediated transformation. HR phenotypes were observed and imaged 2 days post infiltration. To detect protein expressions, Roq1 mutants and StrepII-XopQ were co-expressed in *Nicotiana benthamiana eds1-1* mutant leaves, extracted using protein extraction buffer (50 mM Tris-HCl, pH 7.5, 150 mM NaCl, 1mM EDTA, 0.2% Nonidet P-40, 0.2%Triton X-100, 6 mM β- mercaptoethanol, 10mM DTT and 1×Protease Inhibitor Cocktail), boiled in Laemmli buffer for 5 min and separated on 8% SDS-PAGE gels. StrepII-XopQ were detected with the primary StrepTag II monoclonal antibody (A02230, Abbkine) and the second antibody (A4416, Sigma). Roq1 mutants were enriched by ANTI-FLAG M2 Affinity Gel (A2220, Sigma-Aldrich) before boiled, and detected with the primary monoclonal anti-Flag antibody (F1804, Sigma-Aldrich) and the second antibody (A4416, Sigma).

---

**Fig. S1.**
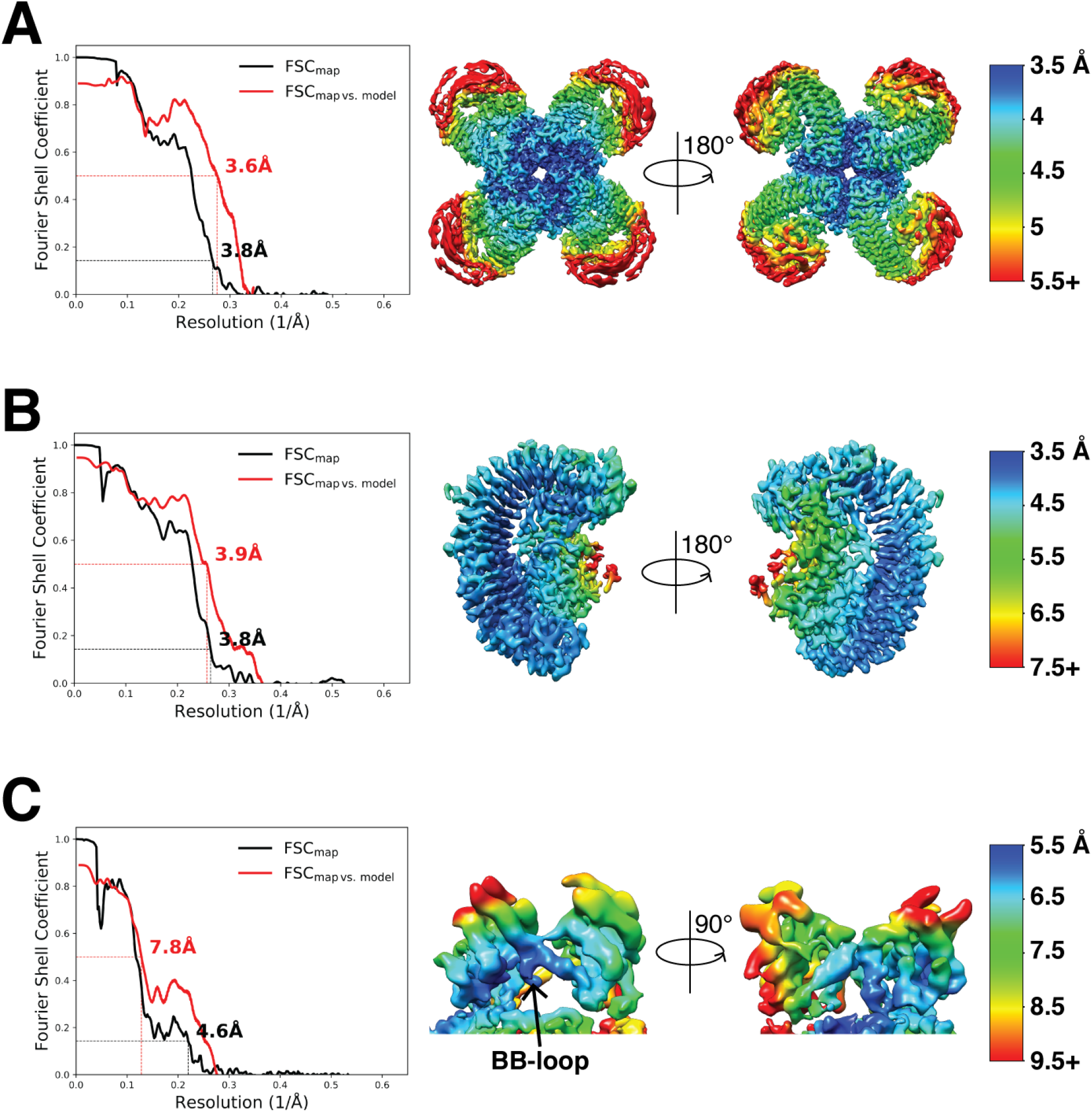
Resolution estimation. Left: map FSC and map vs model FSC for the three reconstructions used to build the atomic model of the Roq1-XopQ complex. Right: Local resolution for each map. (A) Map vs model FSC calculated using residues 189-625 of Roq1, corresponding to the NB-ARC and the N-terminal region of the LRR. (B) Map vs model FSC calculated using residues 526-1303 of Roq1, corresponding to the LRR-PL region, and 89-453 of XopQ. (C) Map vs model FSC calculated using residues 11-176 of Roq1, corresponding to the TIR domain.

**Fig. S2.**
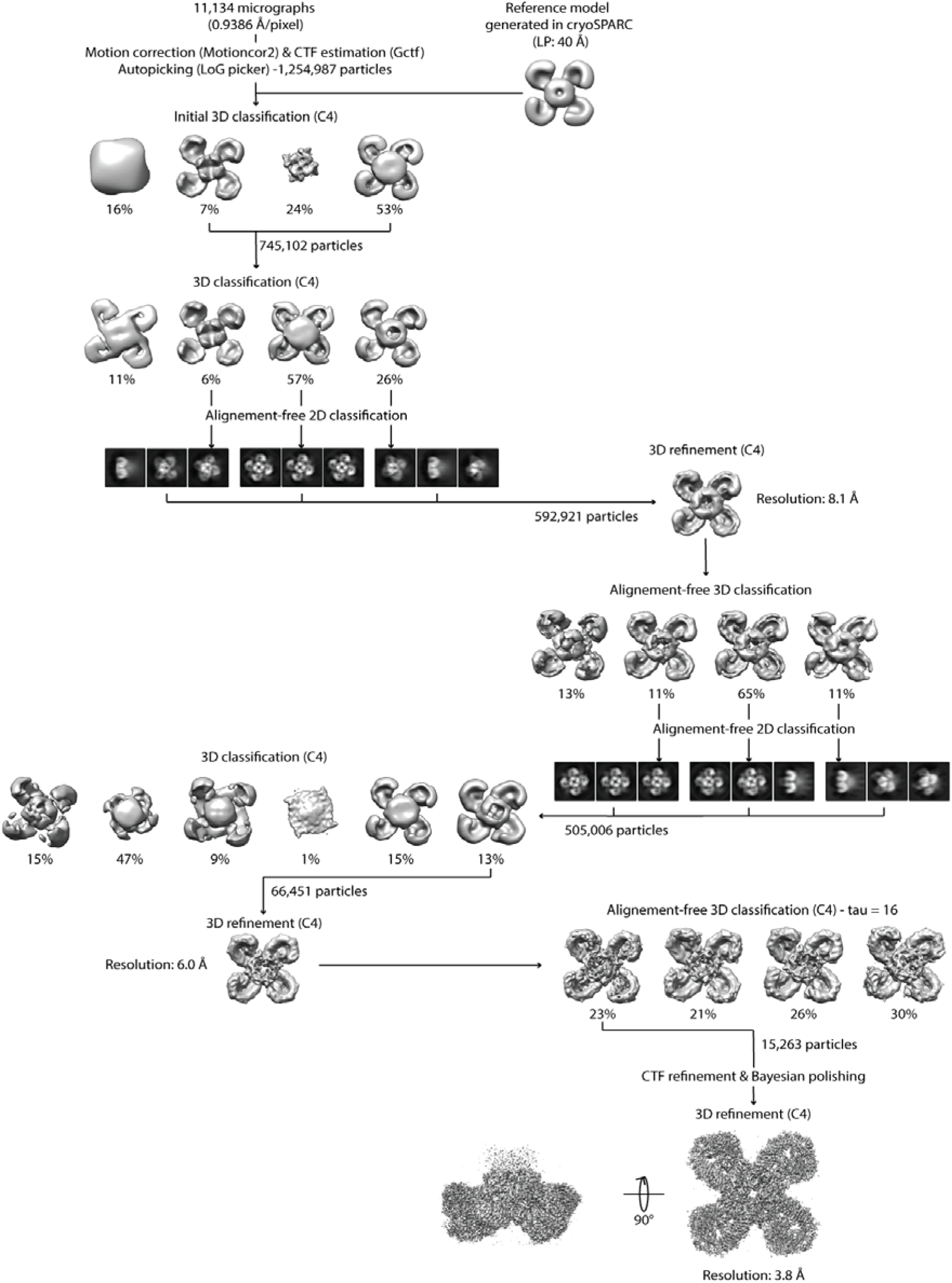
Cryo-EM data processing tree from collected EM movies to the initial reconstruction of the Roq1-XopQ complex.

**Fig. S3.**
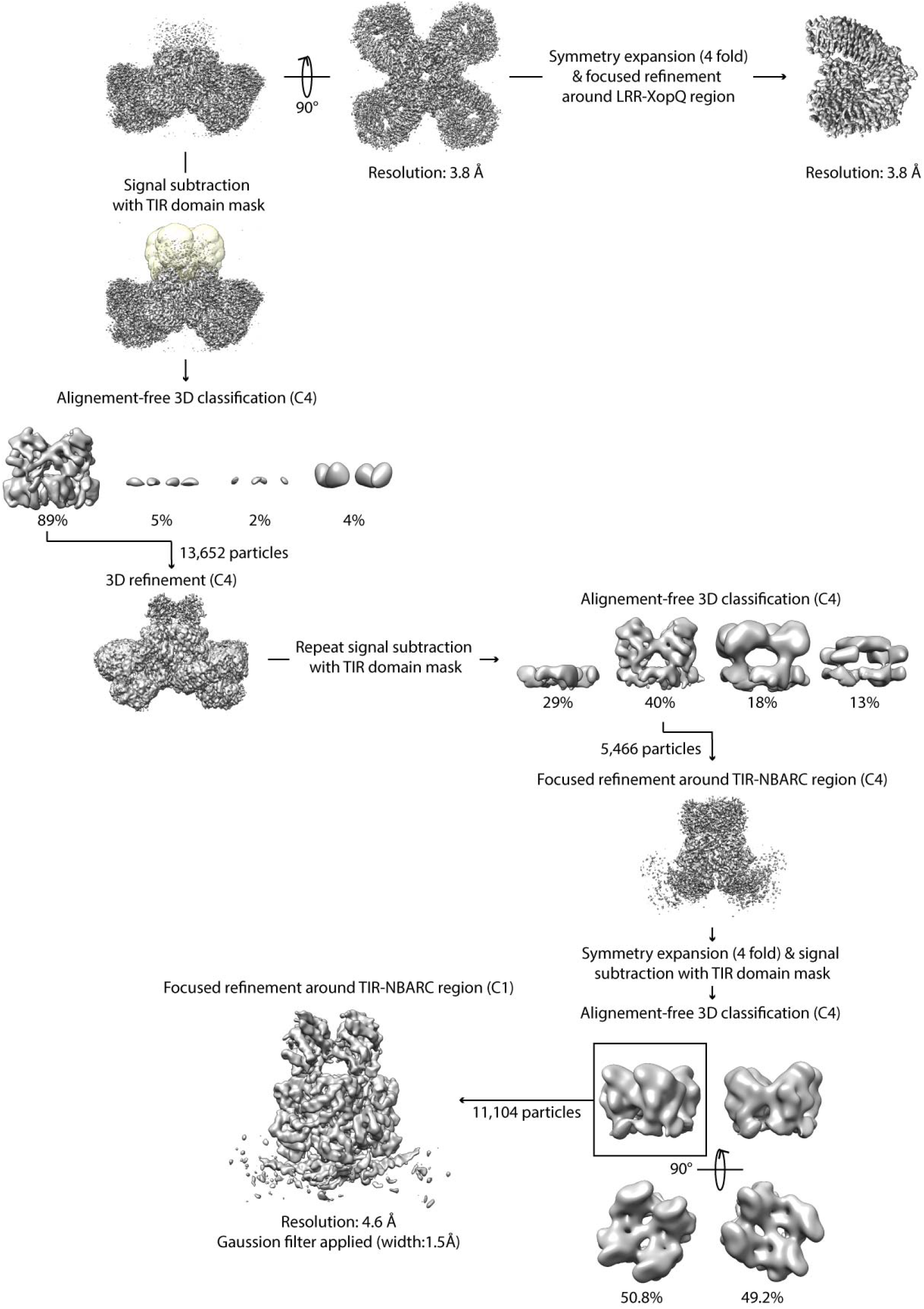
Further cryo-EM data processing needed to resolve the LRR-PL-XopQ region and the TIR domains.

**Fig. S4.**
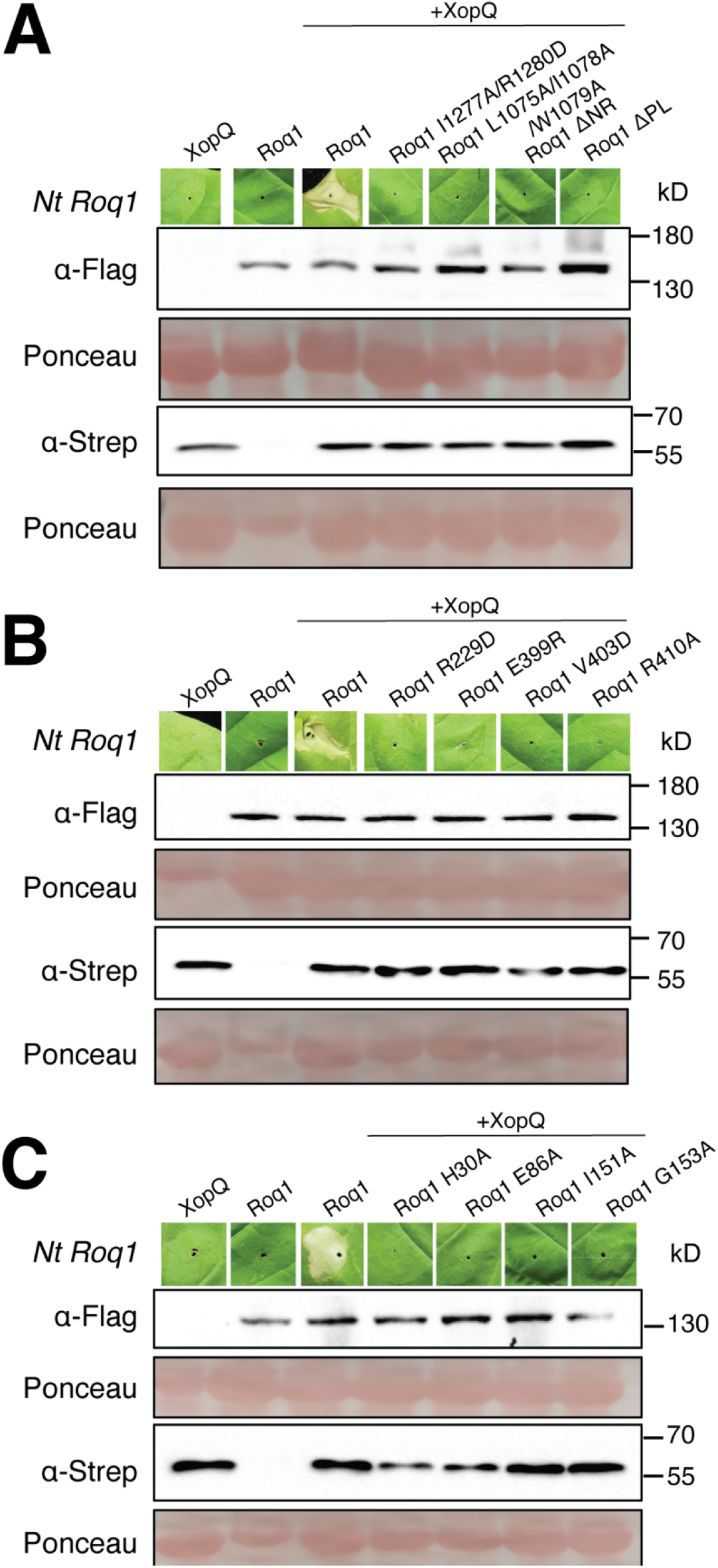
HR Phenotype and Protein Expression of Roq1 mutants. (A) Mutation in the LRR-PL-XopQ region. (B) Mutation in the NB-ARC domain of Roq1. (C) Mutations in the TIR domain of Roq1.

**Fig. S5.**
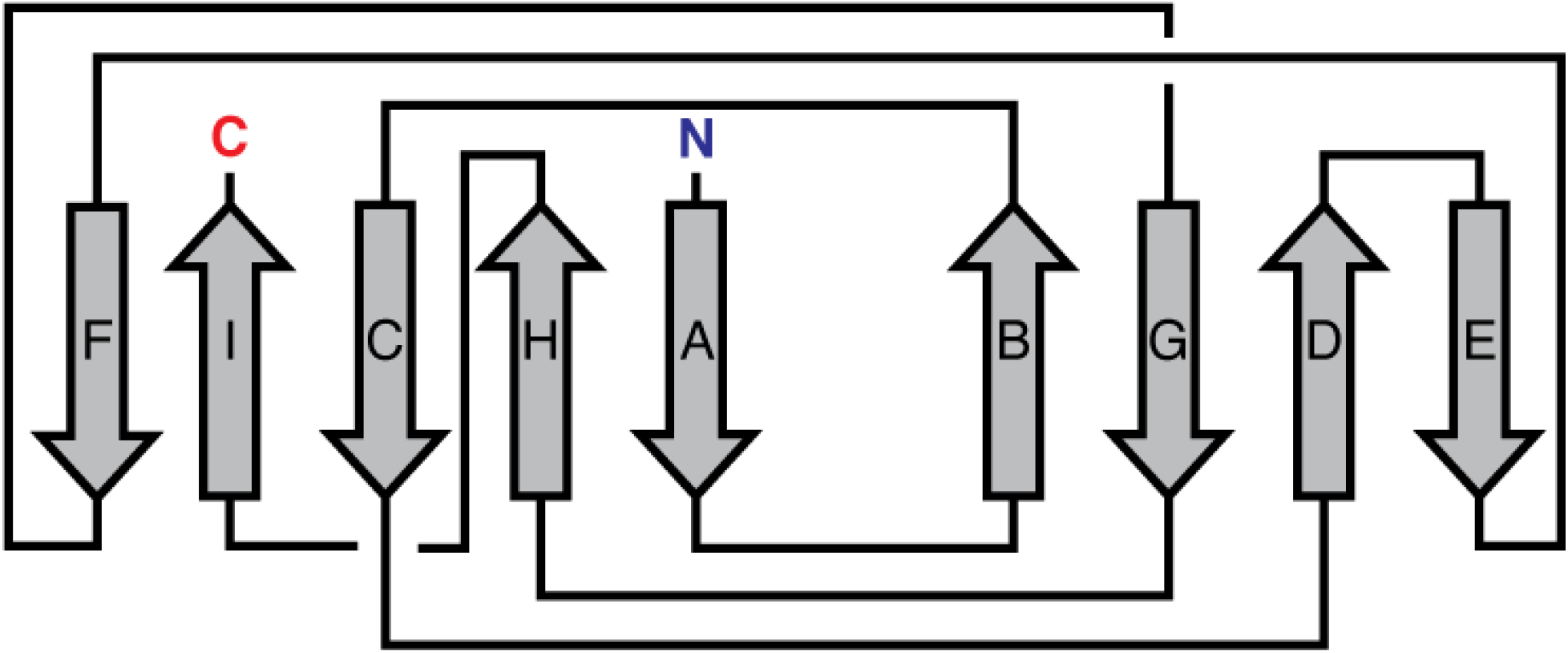
Topology of the PL domain going from N-terminal most β-strand to the C-terminus in alphabetical order.

**Fig. S6.**
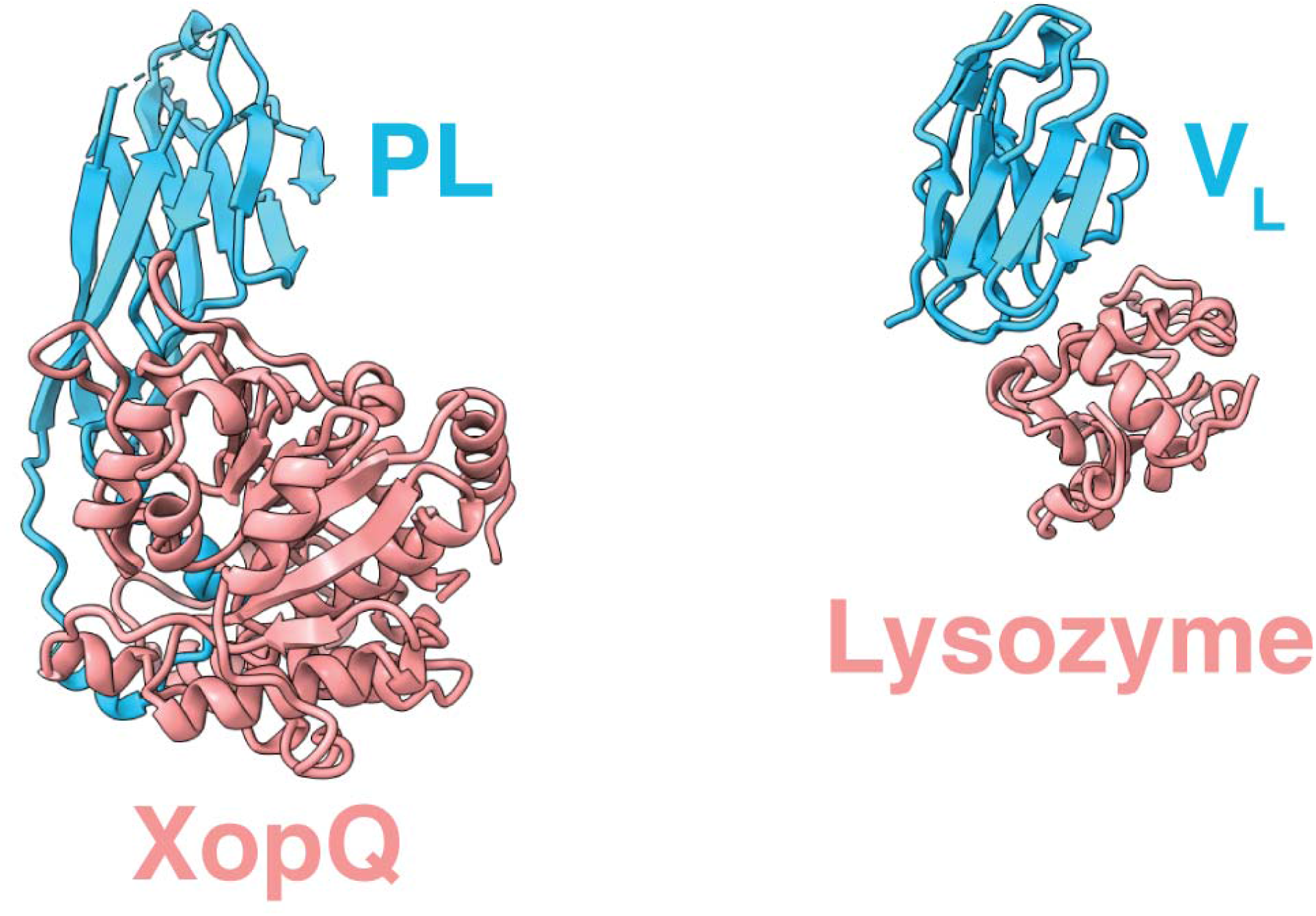
Comparison between the PL domain of Roq1 recognizing XopQ (left) and the light-chain variable fragment (V_L_) of an antibody recognizing lysozyme (PDB: 3HFM) (right). Loops emerging from the β-sandwich of either the PL or V_L_ (light blue) interact with the substrate (salmon).

**Fig. S7.**
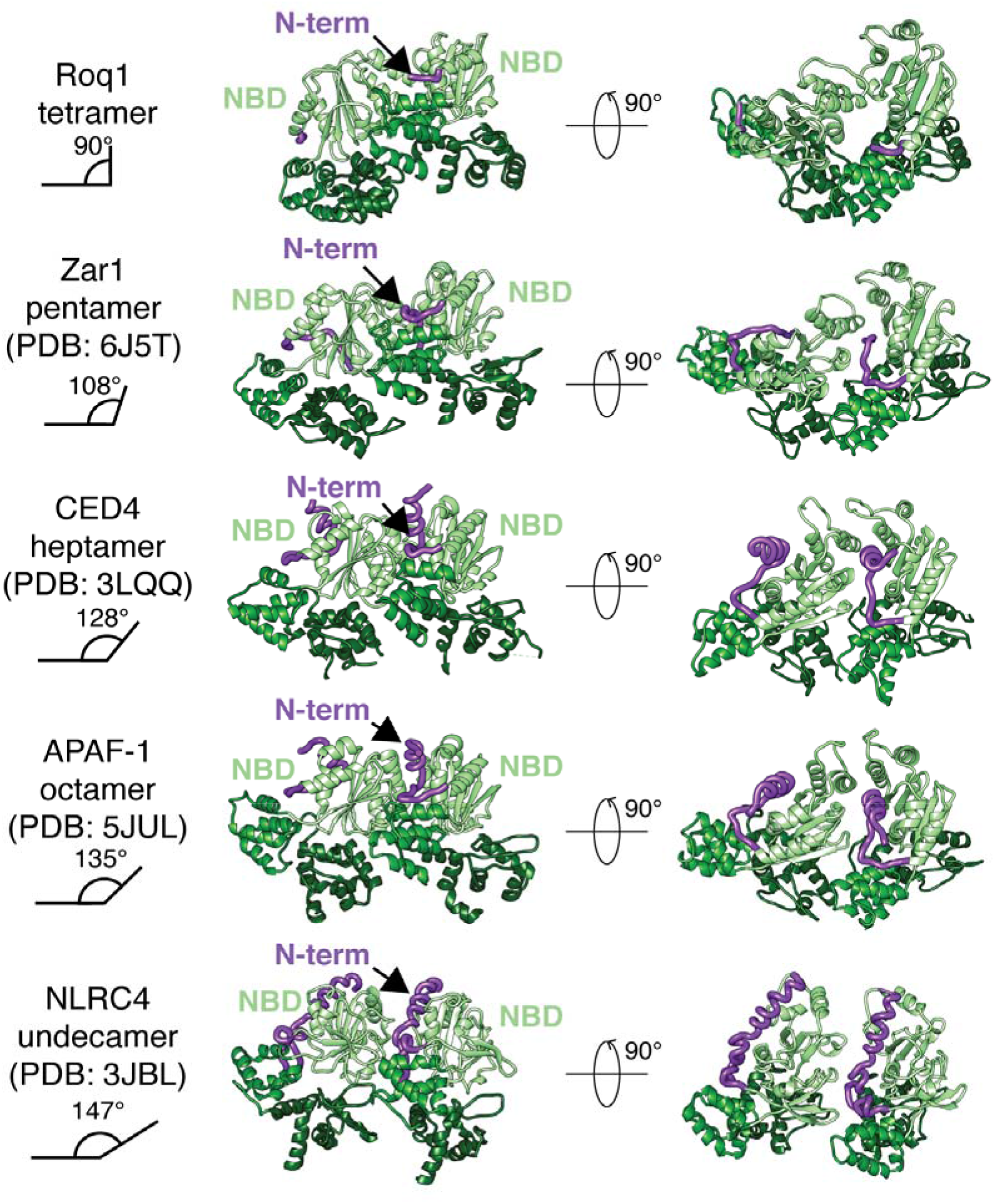
Structure of the NBD N-terminal linker (purple) in activated NLRs with increasing oligomeric states. The NBD-HD1-WHD of two neighboring protomers is shown (following the color scheme of Roq1), with the N-terminal linker of the NBD wedged between them.

**Fig. S8.**
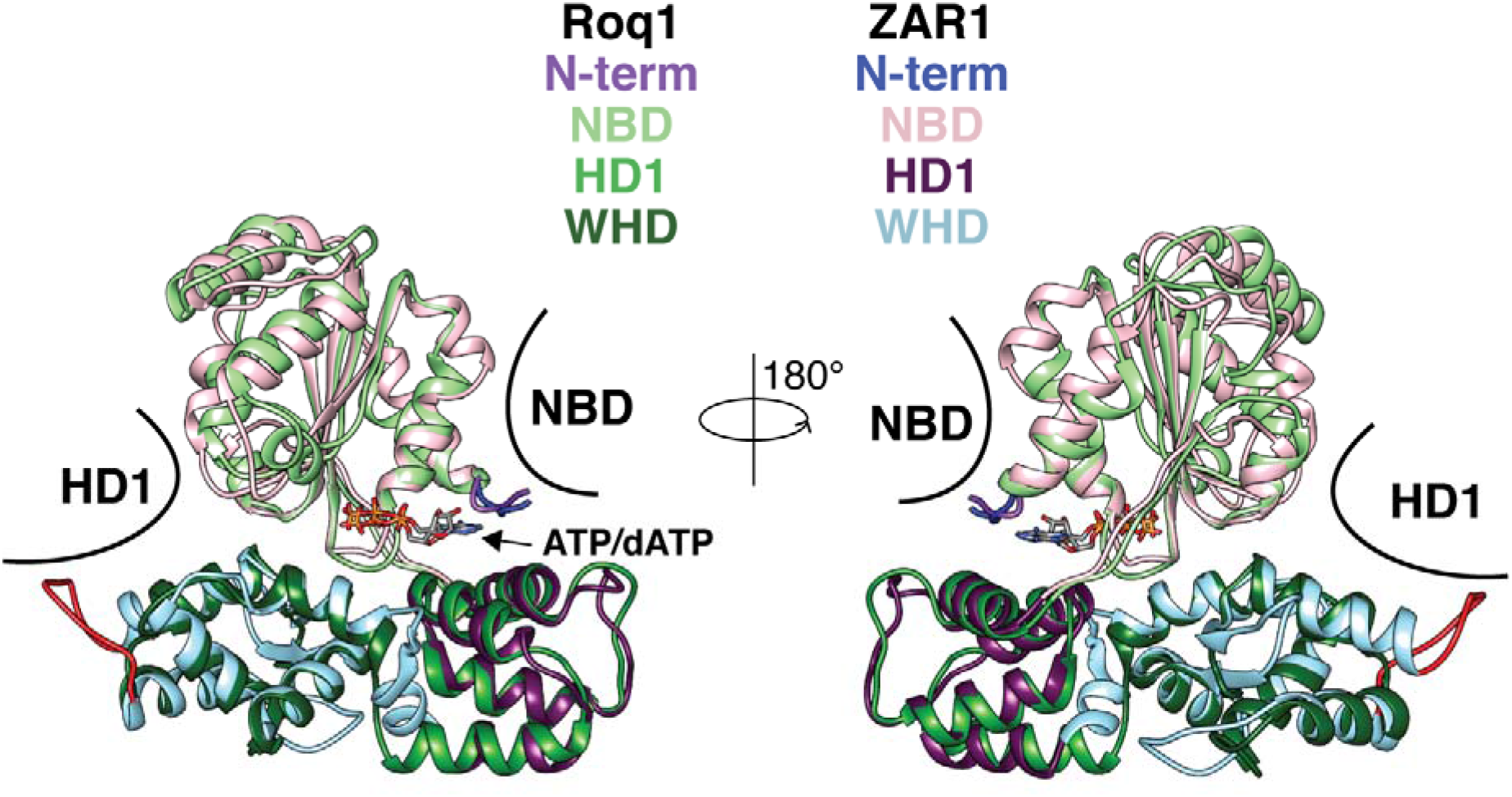
Structural comparison between the NB-ARC domain of Roq1 and ZAR1 in the oligomerized state. The NBD and HD1 of neighboring subunits are represented in black. The NBD and HD1 closely align to each other, whereas we find differences in the loops of the WHD, with one of them extending to make contacts with the neighboring HD1 (red). This loop compensates for the increased distance between the WHD and HD1 in the ZAR1 pentamer relative to the Roq1 tetramer.

**Fig. S9.**
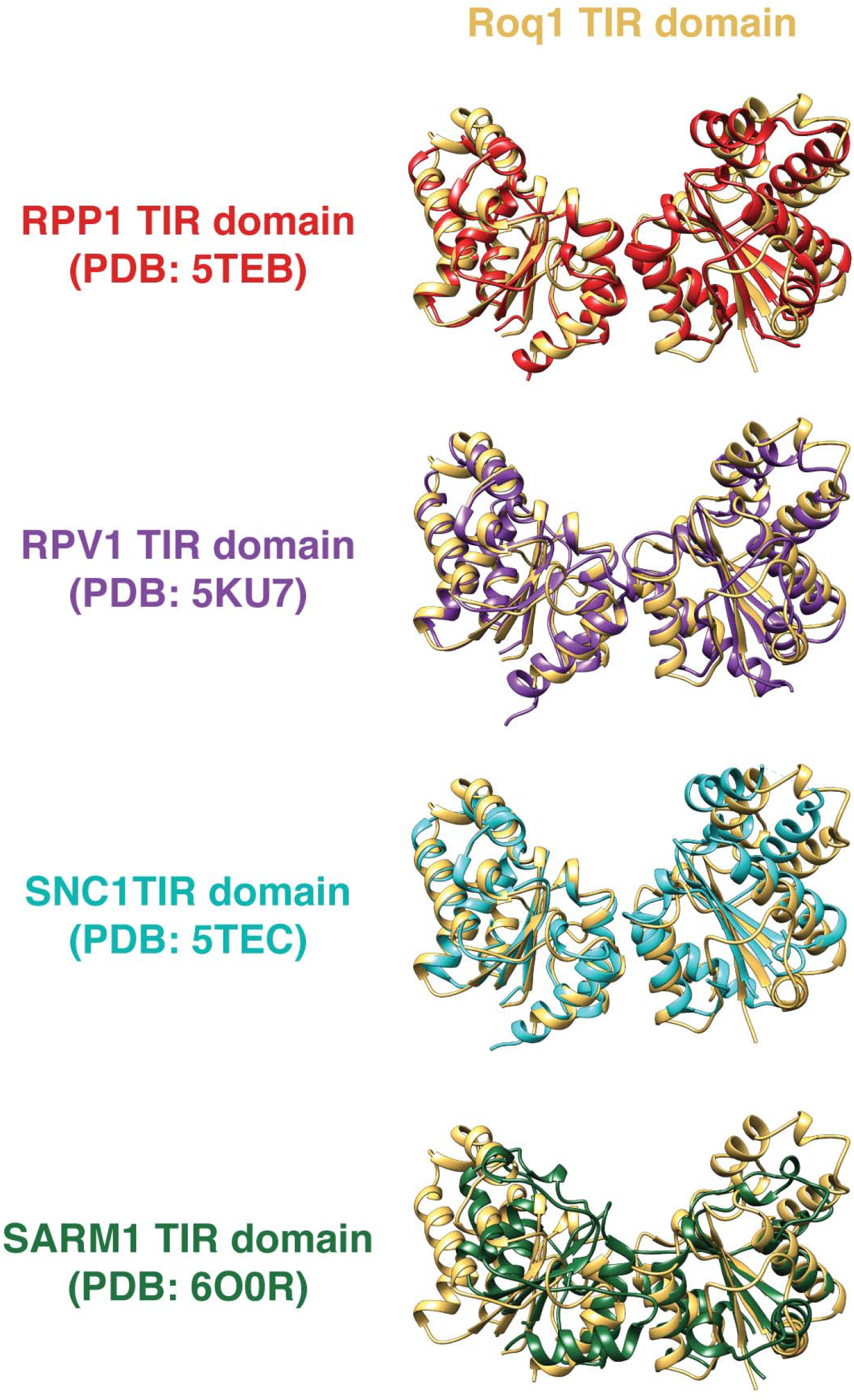
Structural comparison between the Roq1 TIR AE interface and the AE interface found in the crystal lattice of other TIR domains.

**Fig. S10.**
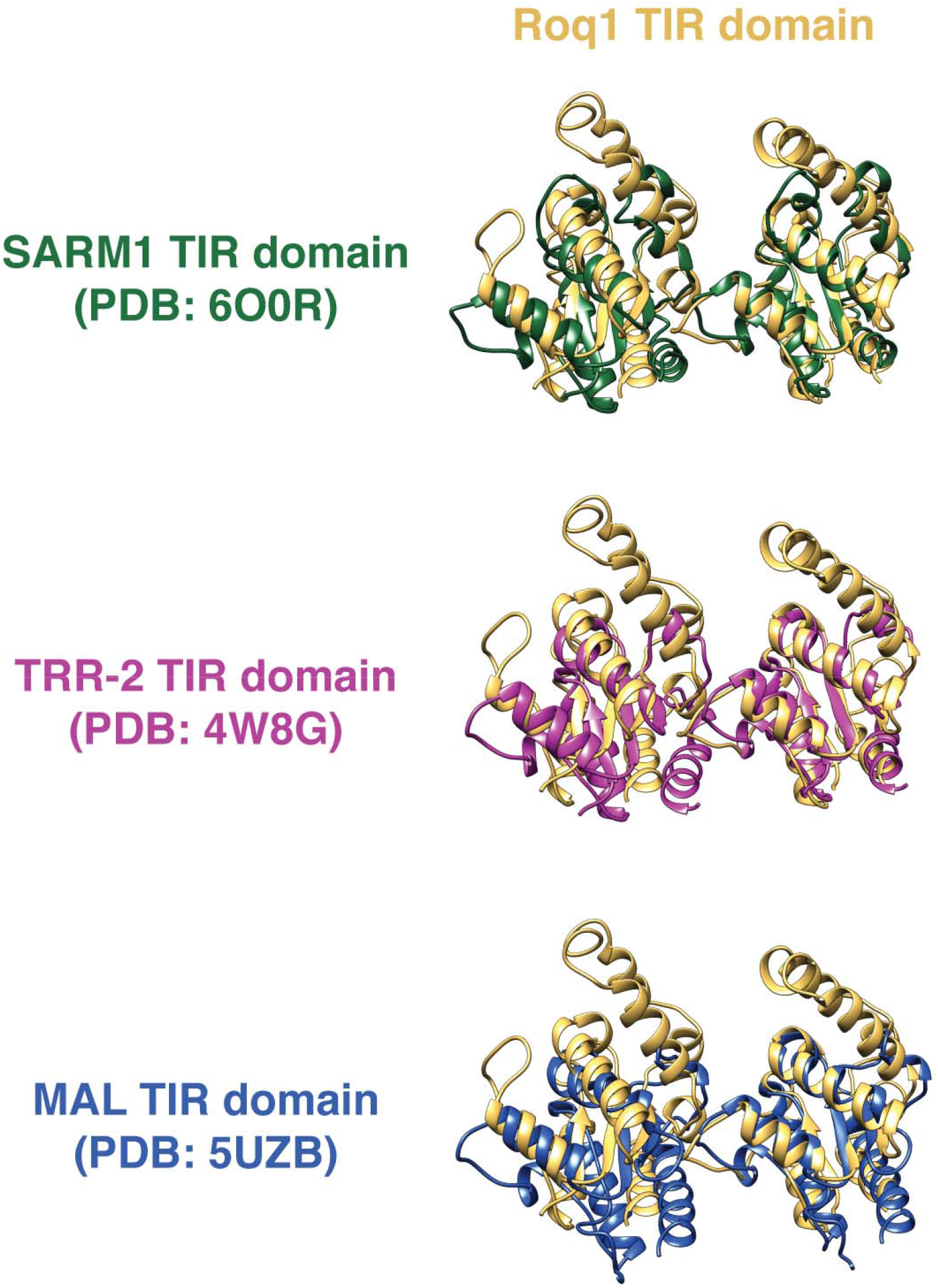
Structural comparison between the Roq1 TIR BE interface and the BE interface found in the crystal lattice of metazoan TIR domains.

**Fig. S11.**
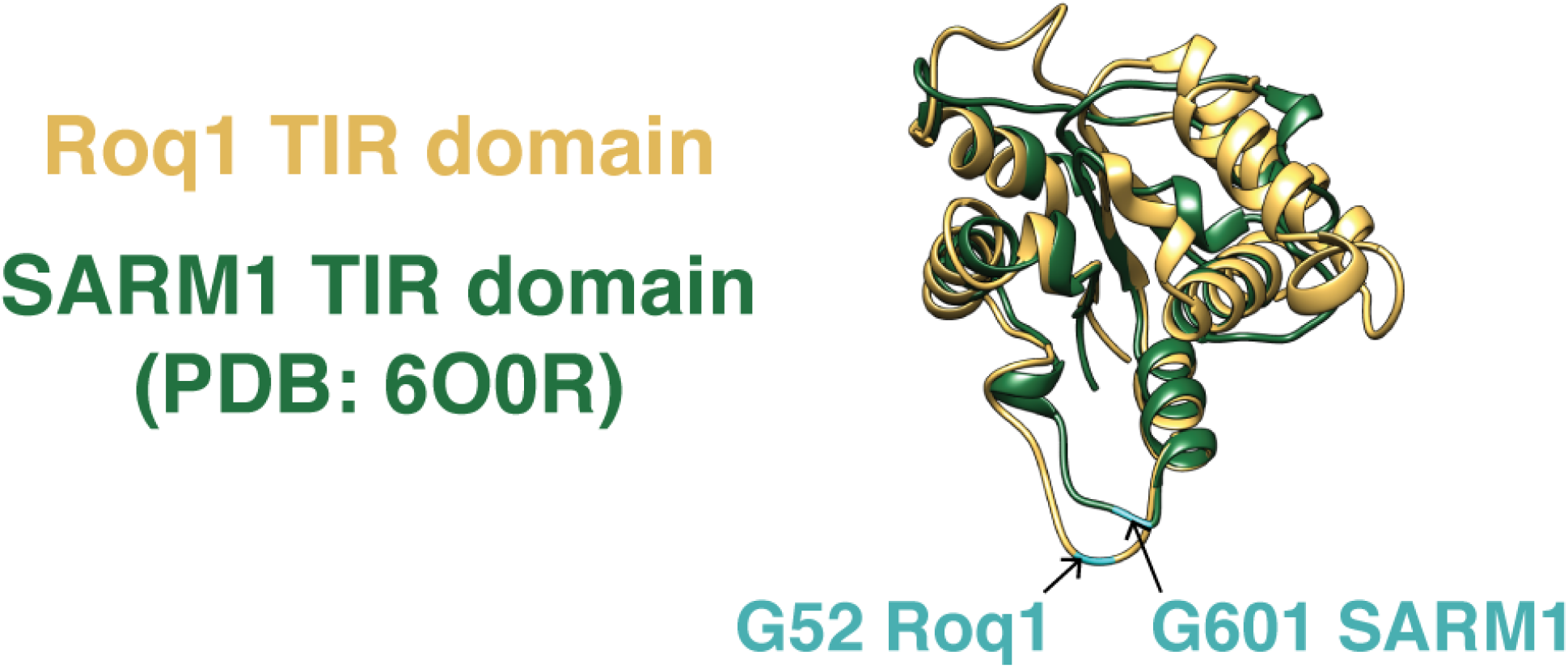
Position of the BB-loop glycine in both Roq1 and SARM1 TIR domains. Glycines are represented in cyan.

**Fig. S12.**
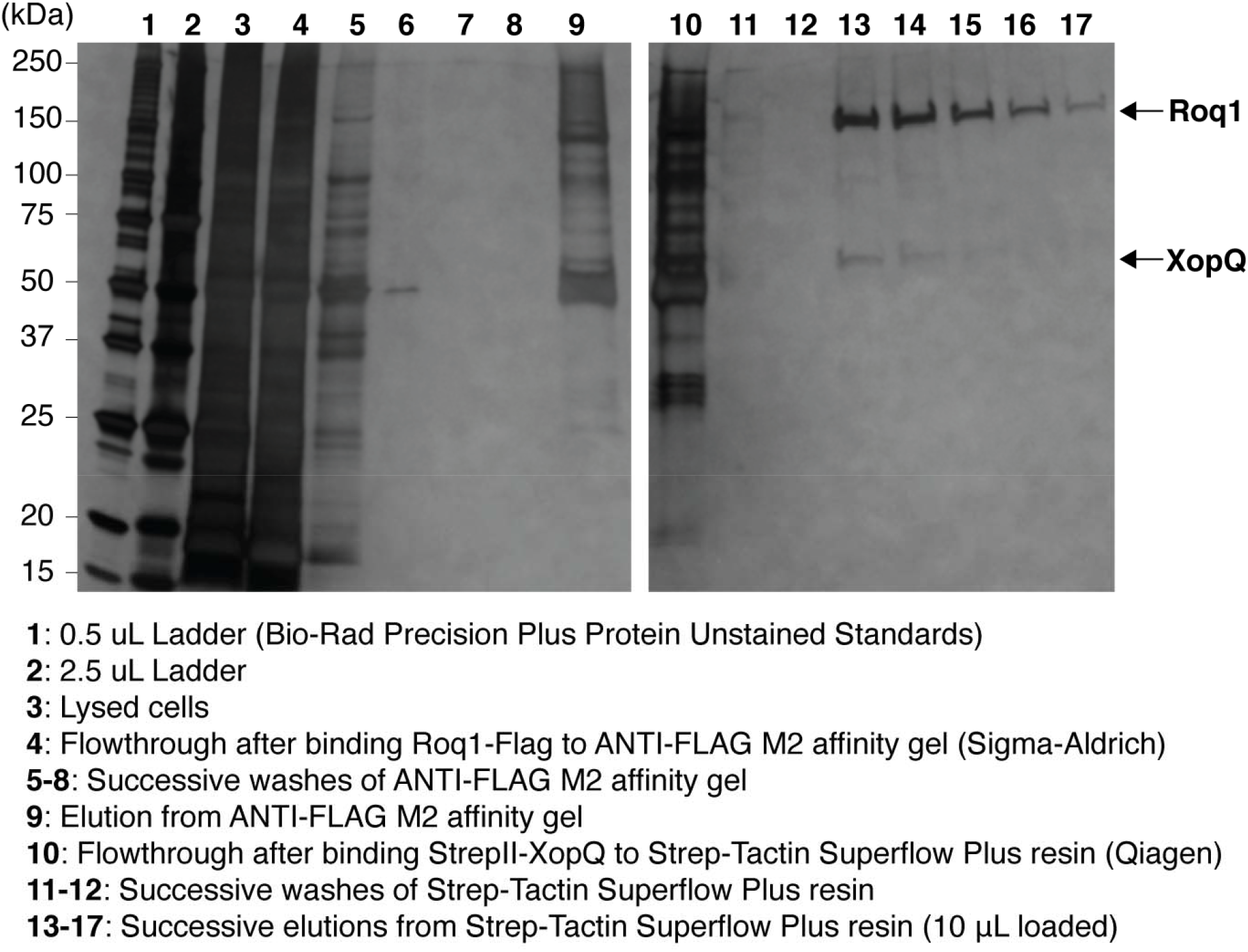
Purification of the Roq1-XopQ complex. Samples were analyzed by SDS-PAGE and silver stained.

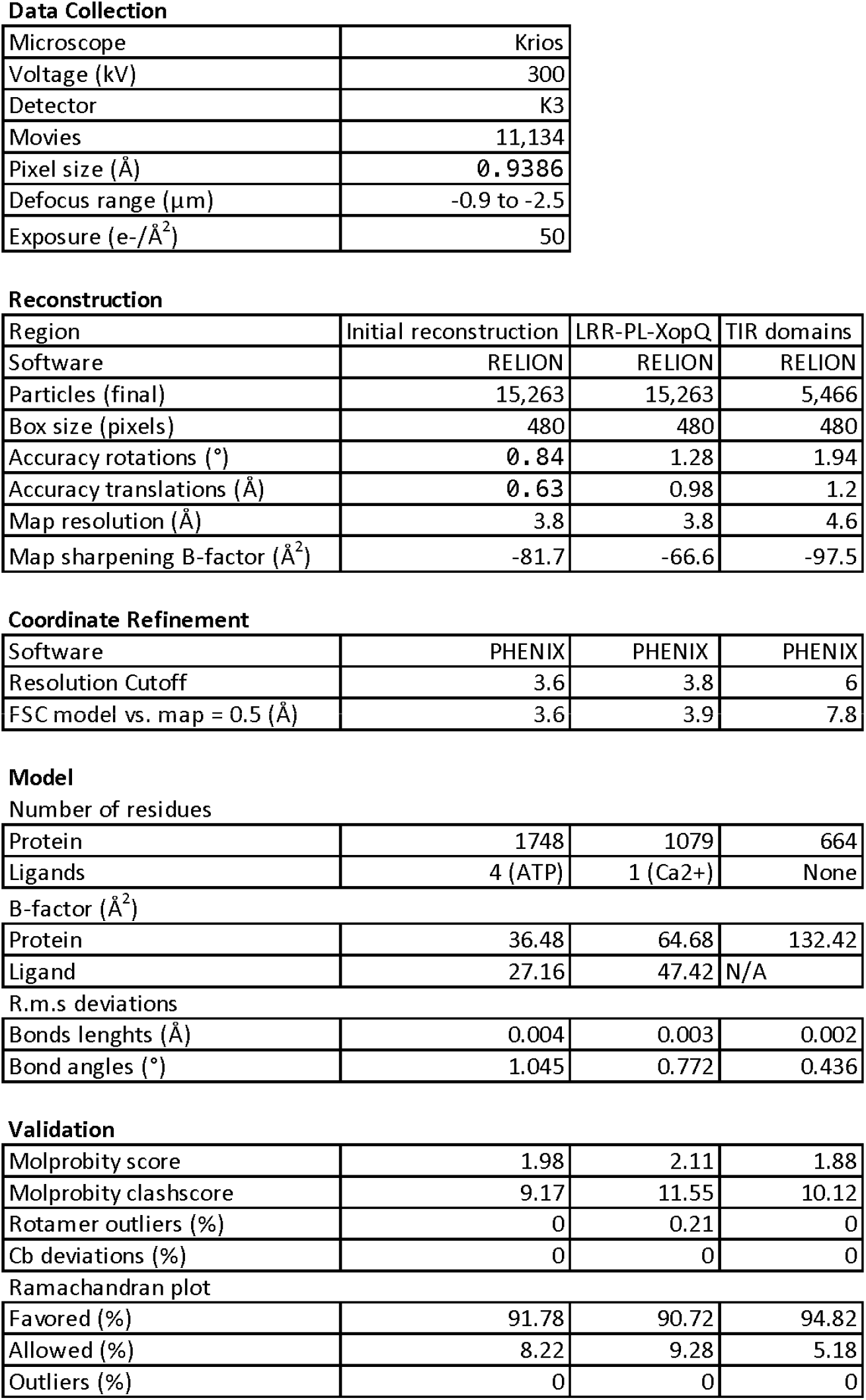
Cryo-EM data collection, data processing, model refinement and validation statistics.

**Data Collection**
|  |  |
| --- | --- |
| Microscope | Krios |
| Voltage (kV) | 300 |
| Detector | K3 |
| Movies | 11,134 |
| Pixel size (Å) | 0.9386 |
| Defocus range (μm) | -0.9 to -2.5 |
| Exposure (e-/Å<sup>2</sup>) | 50 |

**Reconstruction**
| Region | Initial reconstruction | LRR-PL-XopQ | TIR domains |
| --- | --- | --- | --- |
| Software | RELION | RELION | RELION |
| Particles (final) | 15,263 | 15,263 | 5,466 |
| Box size (pixels) | 480 | 480 | 480 |
| Accuracy rotations (°) | 0.84 | 1.28 | 1.94 |
| Accuracy translations (Å) | 0.63 | 0.98 | 1.2 |
| Map resolution (Å) | 3.8 | 3.8 | 4.6 |
| Map sharpening B-factor (Å<sup>2</sup>) | -81.7 | -66.6 | -97.5 |

**Coordinate Refinement**
|  |  |  |  |
| --- | --- | --- | --- |
| Software | PHENIX | PHENIX | PHENIX |
| Resolution Cutoff | 3.6 | 3.8 | 6 |
| FSC model vs. map = 0.5 (Å) | 3.6 | 3.9 | 7.8 |

|  |  |  |  |
| --- | --- | --- | --- |
| Protein | 1748 | 1079 | 664 |
| Ligands | 4 (ATP) | 1 (Ca2+) | None |

B-factor (Å<sup>2</sup>)
|  |  |  |  |
| --- | --- | --- | --- |
| Protein | 36.48 | 64.68 | 132.42 |
| Ligand | 27.16 | 47.42 | N/A |

R.m.s deviations
|  |  |  |  |
| --- | --- | --- | --- |
| Bonds lengths (Å) | 0.004 | 0.003 | 0.002 |
| Bond angles (°) | 1.045 | 0.772 | 0.436 |

**Validation**
|  |  |  |  |
| --- | --- | --- | --- |
| Molprobity score | 1.98 | 2.11 | 1.88 |
| Molprobity clashscore | 9.17 | 11.55 | 10.12 |
| Rotamer outliers (%) | 0 | 0.21 | 0 |
| Cb deviations (%) | 0 | 0 | 0 |

Ramachandran plot
|  |  |  |  |
| --- | --- | --- | --- |
| Favored (%) | 91.78 | 90.72 | 94.82 |
| Allowed (%) | 8.22 | 9.28 | 5.18 |
| Outliers (%) | 0 | 0 | 0 |

**Movie S1.**
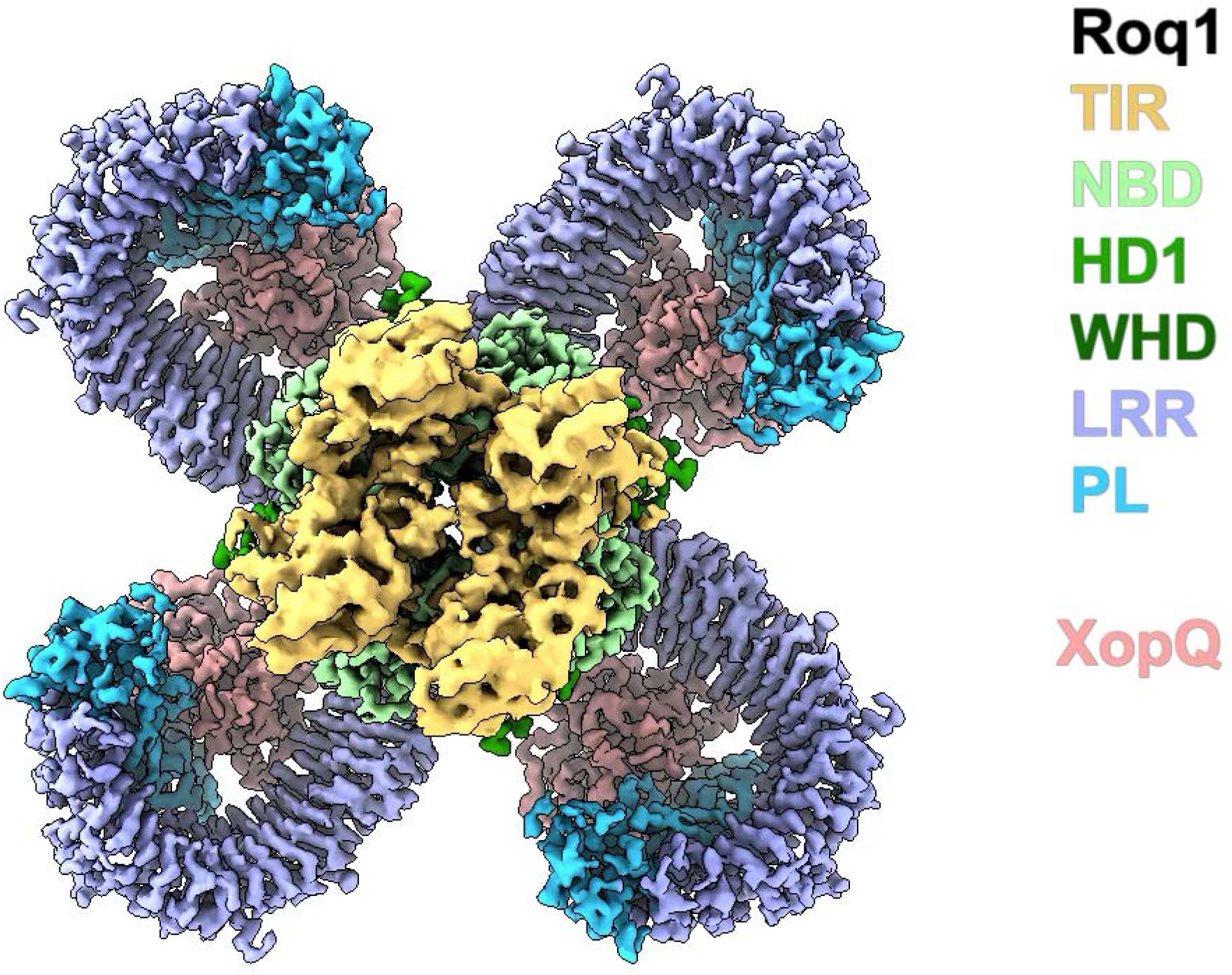
Cryo-EM density map and atomic model of the Roq1-XopQ complex with colors corresponding to the different protein domains (as in Fig. 1). Regions of interest are zoomed in to highlight the different features of the structure.

